# Adhesion and polarity-driven morphogenesis: Mechanisms and constraints in tissue formation

**DOI:** 10.64898/2026.01.23.701437

**Authors:** Yoshiyuki T. Nakamura, Chikara Furusawa, Kunihiko Kaneko

**Affiliations:** Department of Physics, The University of Tokyo, Tokyo, Japan; Universal Biology Institute, The University of Tokyo, Tokyo, Japan; Center for Biosystems Dynamics Research, RIKEN, Kobe, Japan; Niels Bohr Institute, University of Copenhagen, Copenhagen, Denmark

## Abstract

Embryonic development in multicellular organisms exhibits diverse morphogenetic patterns, which can generally be categorized into fundamental types such as monolayer and multilayer spheres, as well as cell masses. Furthermore, we identify two distinct processes for the formation of spherical structures. These basic patterns are thought to be governed by the microscopic properties of intercellular adhesion. However, the specific mechanisms linking the microscopic factors to the emergence of distinct macroscopic morphogenetic patterns remain poorly understood. In this study, we explore how different morphogenetic patterns arise by employing a computational model that incorporates intercellular adhesion and polarity. Our results demonstrate that all fundamental morphogenetic patterns can be generated through the interplay of two key parameters: the strength of cell polarity and the regulation of polarity via mechanical signals. Furthermore, analytical discussions reveal partial mechanisms underlying the formation of these patterns. These findings highlight the critical role of physical constraints in morphogenesis and suggest potential applications in the design principles for artificial tissues and organoids.

**Author summary:** Living organisms build their bodies through morphogenesis, during which cells autonomously arrange themselves into functional structures such as sheets, tubes, and spheres. From simple monolayered spheres to complex multilayered tissues organized by adhesion, it remains unclear how such diverse forms arise. Here, we mathematically modeled a population of proliferating cells governed only by two microscopic factors: the strength of polarity-dependent adhesion and the time scale at which polarity is regulated by cell-cell contact. Surprisingly, we found that this minimal model reproduces five basic morphological types observed in living embryos, including monolayer/multilayer structures and two distinct modes of cavity formation: by wrapping around or by inflating from the inside. Systematic simulations revealed that these macroscale outcomes are determined solely by two parameters controlling polarity strength and its regulation, suggesting that simple physical rules underlie diverse developmental architectures. Analysis of the model uncovers phase transitions between the five morphogenetic types and reveals how varying polarity and adhesion can recapitulate features of real embryogenesis. Our work proposes a unified framework that connects microscopic polarity mechanics to diverse developmental morphologies and provides a foundation for future applications in organoid design and tissue engineering.

## 1 Introduction

The development of multicellular organisms begins with a single cell, which proliferates and organizes into diverse macroscopic structures. Despite this shared starting point, the manner in which early embryonic cells arrange themselves varies significantly across different species. In zebrafish embryos, multiple cell layers form as the blastoderm spreads over the large yolk mass, a process characteristic of discoidal cleavage (Fig. 1) [1–3]. In amphibian embryos, cells generate a multi-layered sphere by inflating from within, with differing cell layer numbers between the animal and vegetal poles (Fig. 1) [1, 4]. Mammalian embryos initially form a compact solid cluster, which later reorganizes into a blastocyst consisting of an outer monolayered epithelium and an inner cell mass [1, 5]. Echinoderm embryos, by contrast, develop into a monolayered hollow sphere (Fig. 1) [1, 6]. Notably, even in choanoflagellates, the closest unicellular relatives of animals, cells form a monolayered sphere, involving the closure of an initially arc-shaped configuration (Fig. 1) [7, 8].

**Fig 1.**
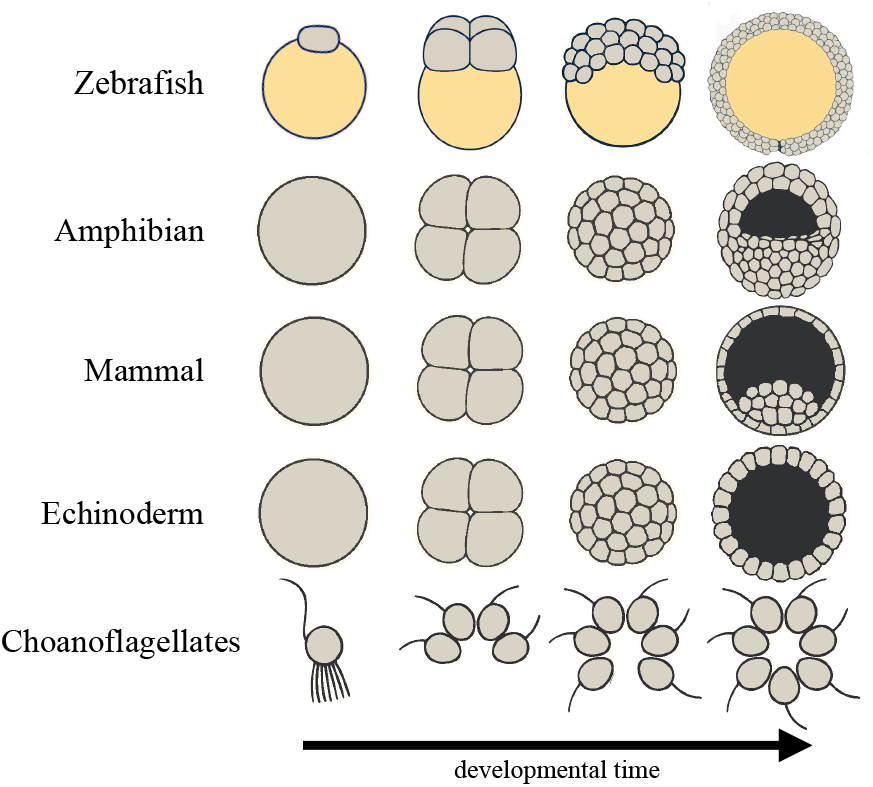
Illustrated comparison of early embryogenesis across metazoan lineages and colony formation in choanoflagellates. Gray regions represent cells, while yellow regions indicate yolk. The horizontal axis represents developmental time.

As observed in a variety of early embryos, we note that there are basic typical behaviors: cells form either masses or hollow spheres. When forming a sphere-like structure, the cell layer on the surface can be either single-layered or multi-layered in which the number of layers can vary. Additionally, in the case of sphere formation, there are two types: either a sphere is formed by wrapping around or it is formed by inflating from the center core cells. Thus the behaviors can be categorized into 2 v2 classes of mono/multi-layered and wraparound/inflation, in addition to the mass formation case (Fig. 2). We refer to these 5 classes as the “basic types.”

**Fig 2.**
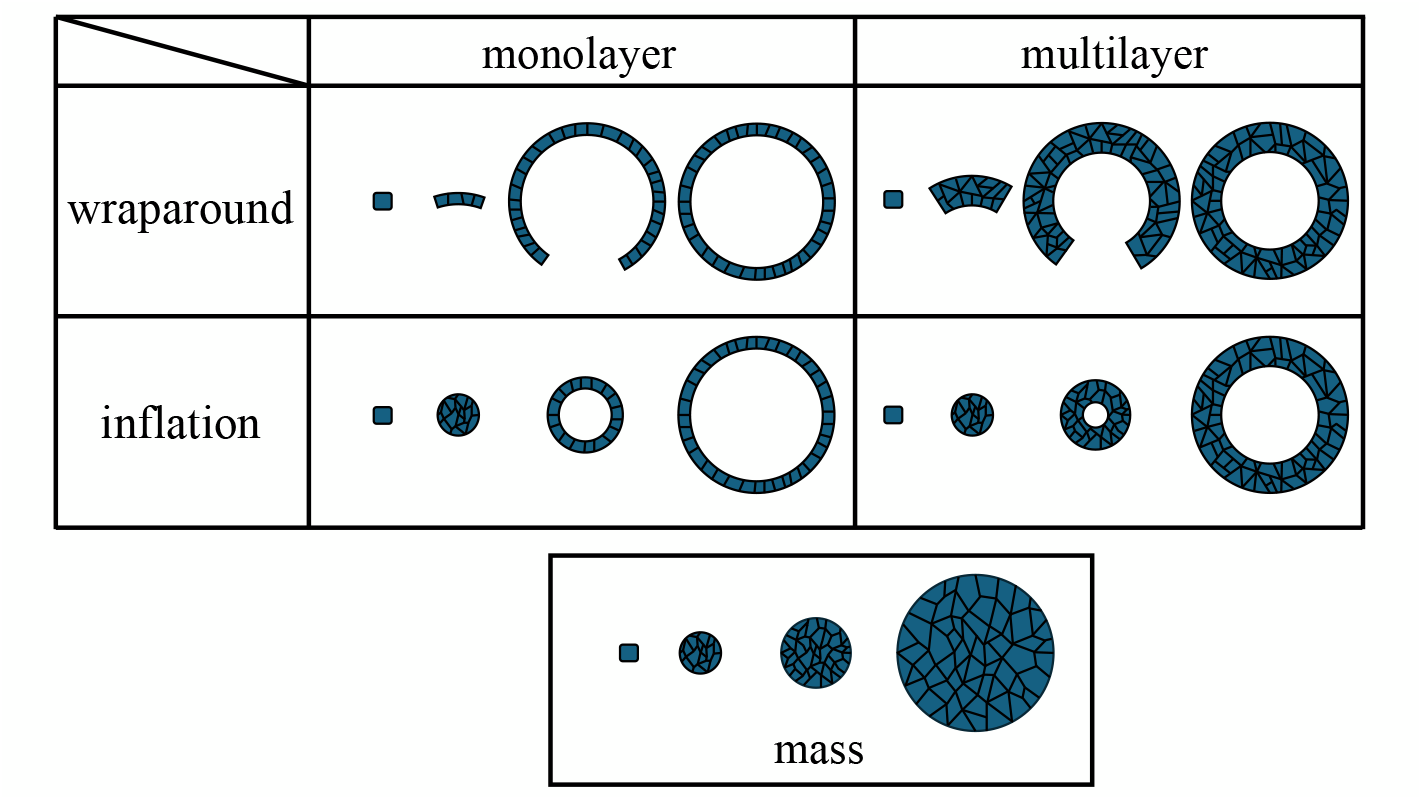
Categorization of the 5 basic types. The horizontal axis represents developmental time.

The diversity in morphogenetic patterns among these organisms does not appear to correlate with their phylogenetic relationships. This suggests that the mechanisms responsible for their differences may not be solely determined by genetic relatedness. Can we understand the basic types of morphogenesis as generic physical mechanisms [9, 10]? What mechanisms contribute to generating distinct morphogenetic processes? One of the main drivers of morphogenesis is the sequential arrangement of individual cells while maintaining adhesion [9]. Cell adhesion is primarily mediated by adhesion proteins, with cadherins being a well-known example, which are a class of transmembrane proteins that facilitate cell-cell adhesion by forming adherens junctions [11]. Cadherins interact with catenins, which link the cadherins to the actin cytoskeleton, thereby providing mechanical stability and signaling functions [12]. The differential expression of cadherins between cell types can lead to the segregation of cell populations and the formation of distinct tissue structures [13]. For example, E-cadherin is crucial for the formation of epithelial layers, while N-cadherin is involved in the development of neural tissues [14].

Another important factor in cell-cell interaction is apico-basal polarity, which plays a crucial role in determining the mode of adhesion and, consequently, morphogenesis [15, 16]. Apico-basal polarity refers to the spatial differences in the distribution of cellular components along the apico-basal axis of epithelial cells. This polarity is essential for the formation of proper epithelial tissues, as it is a dominant factor in determining the orientation of cells and formation of distinct cellular domains. The establishment and maintenance of apico-basal polarity are regulated by a complex network of polarity proteins, including the PAR complex, which interacts with cell adhesion molecules to coordinate cell positioning and tissue architecture [17]. Importantly, cell-cell adhesion is crucial for the establishment of apico-basal polarity by organizing the spatial distribution of polarity proteins, microtubules, and actin filaments, and coordinating their interactions with the cytoskeleton and cell membrane components [18, 19]. Therefore, understanding the interplay between cell adhesion and polarity is essential for elucidating the mechanisms of morphogenesis.

Several theoretical models of morphogenesis that consider the mechanical effects of polarity and adhesion have been proposed. For example, there are models that recapitulate the formation of a monolayer sphere by adding polarity to the cellular Potts model [20–22] or the cellular vertex model [23–26]. Additionally, cell-based models including polarity-dependent adhesion are also used to discuss the origin of the complexity of shapes [27] and robustness [28], from which the influence of polarity on the formation of morphology has been investigated.

Of course, the actual morphogenesis involves complicated processes. Still, it will be important to explore how much of morphogenesis, in particular the five basic types, can be understood just by the adhesion and polarity. So far, however, consequences of these two factors for macroscopic morphogenetic patterns are poorly understood. In this study, we investigate the potential outcomes that emerge when considering only adhesion and polarity as the first step toward understanding the difference in morphogenesis among species. Following the spirit of statistical physics that links differences in macroscopic phases to variations in microscopic properties, we classify the basic types of morphogenesis that arise as distinct “phases” from differences in cell-associated parameters. Specifically, variations in cell polarity and adhesion strength may serve as critical determinants of these types. Furthermore, it aims to examine the mechanism of how the differences in microscopic factors give rise to different basic types of morphology. This approach provides a framework to connect micro-level cellular parameters with macro-level developmental outcomes, offering novel insights into the principles underlying tissue organization.

Here, for simplicity, we consider possible patterns of morphology formed only by adhesion among cells in a homogeneous population without given specific boundary conditions. Our model consists of a population of cells that proliferate while following the potential for adhesion and the dynamics of polarity control. We do not consider cell differentiation, to focus on only the physical force by adhesion and polarity, as a first step to understand the basic types of morphogenesis. By changing the strength of the polarity and the regulation of the polarity, we study how the morphogenesis process is altered. In addition, we uncover why the number of layers and the formation process differ by using simple analytical calculation.

From our results, we suggest that the number of layers and the formation process can be explained by the difference in adhesion. This result may also be applied to the design of organoids and other artificial tissues.

## 2 Model

We propose a model for the morphogenesis of a cell population. In this model, cells proliferate while following the potential for polarity-dependent adhesion and the dynamics of polarity control. Regarding cell polarity, we consider only one type here, assuming the apico-basal polarity as in epithelial cells. It should be noted that other types of polarity, such as planar cell polarity, are not considered here.

In addition, we primarily consider the case of two-dimensional space. The model can be extended to three dimensions, as discussed in Section 3.4.

### 2.1 The dynamics of cell position and polarity

Each cell has polarity, which is represented as a vector of magnitude *p* (Fig. 3a).

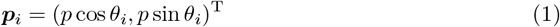

**Fig 3.**
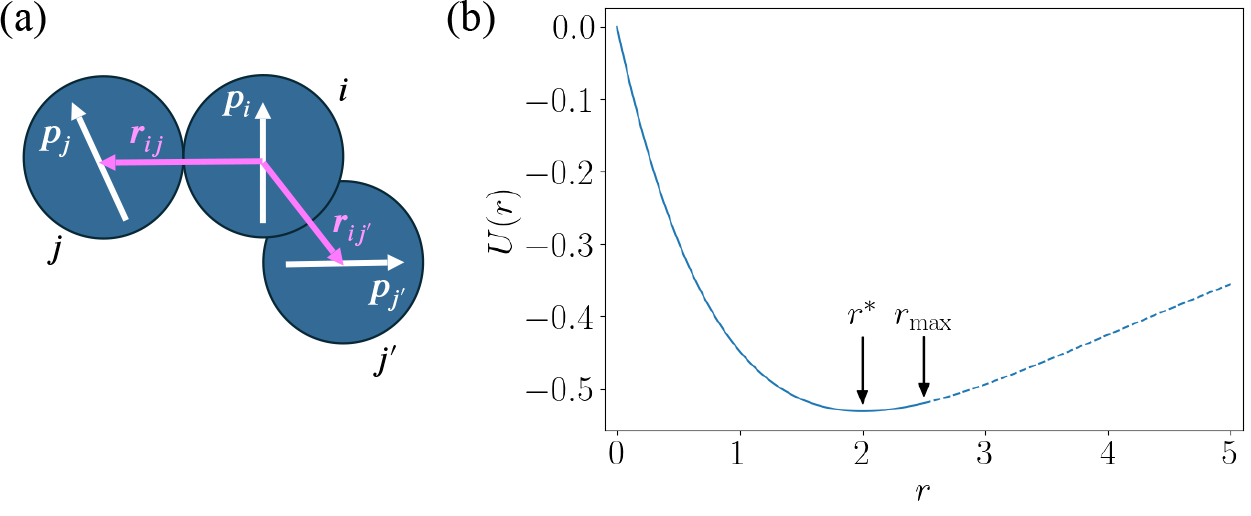
(a) Schematic representation of the model. Cells are represented as blue circles. All cells have the same typical radius *r*^∗^/2, which is set to be 1. Each cell has an intrinsic polarity vector (white arrows). Cells move while interacting with each other. Their dynamics depend on both polarity and the relative position. (b) The shape of the function *U* (*r*). It takes a minimum at *r*^∗^ = 2 (i.e., the radius of the cell is 1). In the region where *r > r*_max_ = 2.5, there is no interaction, which is indicated by the dotted line.

In this model, two cells interact with each other through adhesion. The adhesion is given by short-ranged interaction where there is a cutoff distance of the interaction, which will be discussed later. When two cells interact, they follow dynamics which depend on those positions and polarities. Representing the positions of cells *i* and *j* by ***r***_*i*_ and ***r***_*j*_ respectively, and the distance between them by *r*_*ij*_, the potential *V*_*ij*_ for adhesion force between cells *i* and *j* is described as

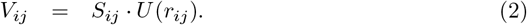

Here, *S* is a factor dependent on polarity, and *U* is a distance-dependent function, which are given by

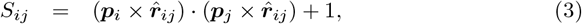

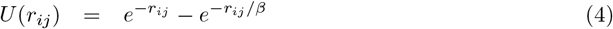

where ***r***_*ij*_ = ***r***_*j*_ −***r***_*i*_ and 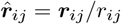. As can be seen from the equations, *S*_*ij*_ is maximized when the polarity of cell *i* and that of cell *j* are in the same direction and perpendicular to 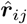. *U* (*r*_*ij*_) is always negative and is minimized when the distance between cell *i* and cell *j* satisfies *U ^′^*(*r*^∗^) = 0. It should be noted that since the value of *r*^∗^ does not depend on the value of *S*_*ij*_, all cells in this model can be approximated as circles with a radius of *r*^∗^/2. For *r < r*^∗^, d*U/*d*r* < 0, so that there is a repulsive force between cells, and for *r* > *r*^∗^, d*U*/d*r >* 0, so that there is an attractive force towards the equilibrium distance *r*^∗^. This force should be short-ranged, and here, we introduced a cutoff distance *r*_max_ beyond which *U ^′^*(*r*) = 0, as will be shown later.

In lowering this potential value *V*_*ij*_, the polarity is aligned due to the effect of *S*_*ij*_, and the radius of the cell approaches *r*^∗^/2 due to the effect of *U* . Here, we determine *β* so that the radius of the cell is 1, that is *r*^∗^ = 2.

As mentioned before, we set the maximum distance of interaction *r*_max_ to consider only local interactions (Fig. 3b). In other words, when *r*_*ij*_ exceeds *r*_max_, the interaction between *i* and *j* does not work. Let *N*_*i*_ be the set of *j*( *i*) such that *r*_*ij*_ < *r*_max_, then the potential *V*_*i*_ applied to cell *i* is the sum of interactions with other cells *j* ∈ *N*_*i*_.

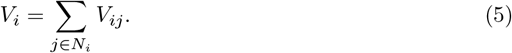

Here, we set *r*_max_ = 2.5.

The position ***r***_*i*_ follows the overdamped dynamics using the above potential.

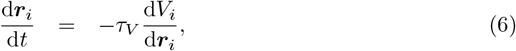

where *τ*_*V*_ is a parameter that represents the time scale of the velocity.

On the other hand, the polarity *θ*_*i*_ is controlled not only by the potential *V*_*i*_ but also by adhesion. Here we choose the dynamics

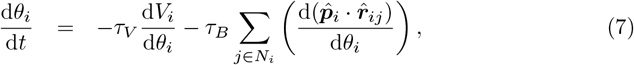

where the second term represents the polarity control with the time constant *τ*_*B*_. Here, 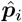 is the unit vector of ***p***_*i*_. With this control, the polarity is directed so that the dot product of the position 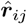 and polarity ***p***_*i*_ can be minimized. In other words, the polarity of cell *i* is forced to face the opposite direction to the adhesive surface. This reflects the biological observation that the polarity is controlled by adhesion or mechanical stress in the actual cell. Several experimental results support the fact that apico-basal polarity is controlled by mechanical stress and is often oriented perpendicular to the adhesion surface [18, 19]. The validity of this assumption will be discussed further in the discussion section.

### 2.2 Cell division

In this model, we consider the cell division process to increase the number of the cells. This process is implemented by randomly selecting a single cell at a certain fixed interval (*τ*_div_) needed for cell division and dividing it. The new cell is generated at a position adjacent to the parent cell. That is, when a cell *i* divides, a new cell *i*^′^ is randomly placed on the circumference with a radius of *r*^∗^ centered on the position ***r***_*i*_ of cell *i*. The polarity of the new cell *i*^′^ has a random angle *θ*^′^. The daughter cell is initially placed at a random position adjacent to the parent cell. Then, its position and polarity subsequently evolve under the forces described by Eqs. (6) and (7), leading the daughter cell to quickly relax into an appropriate configuration.

In this study, we focus on morphogenesis driven by polarity-dependent adhesion. In the cell division method introduced above, a cell division process that is too fast compared to the dynamics of position and polarity is not considered. If the division is too fast, it always dominates over the effect of polarity. In fact, such rapid division just leads to the formation of a mass regardless of polarity or unstable phenomena such as buckling. These phenomena are beyond the scope of this study. We will revisit the validity of this method in the discussion section. We will also discuss the issue of placement of daughter cells in the discussion section.

## 3 Results

The essential dynamics of our model arise from the interplay between adhesion and polarity. The strength of polarity, controlled by *p*, determines how rigidly neighboring cells maintain their relative positions. In addition, the polarity vectors tend to orient in a mutually repulsive manner; the ratio *τ*_*B*_/*τ*_*V*_ governs this local polarity arrangement. This local polarity arrangement, in turn, constrains the emergence of global morphology. In the present analysis, we fix *τ*_*V*_ and vary *τ*_*B*_ to explore how polarity affects morphogenesis.

With this setup, we first show that the model can reproduce the types of morphologies introduced in the introduction by tuning *p* and *τ*_*B*_. We then provide an analytical estimate of the phase boundaries, followed by an extension of the model to generate more complex morphologies. Finally, we demonstrate that the model can be generalized to three dimensions, which does not substantially alter the two-dimensional results.

### 3.1 Typical behaviors

First, we present some representative behaviors, which appear depending on the system parameters *p* and *τ*_*B*_ (See Fig. 4). Note that all the results shown below are calculated up to *t* = 3 × 10^5^, and *τ*_div_ = 10^3^. The initial cell is placed at the center of a square region with a width of 100 in both *x* and *y* axes. We adopt boundary conditions where each cell is confined within this square region. However, if even a single cell comes into contact with the edges of the region, it is not used in subsequent analyses, so these boundary conditions do not affect the main results.

**Fig 4.**
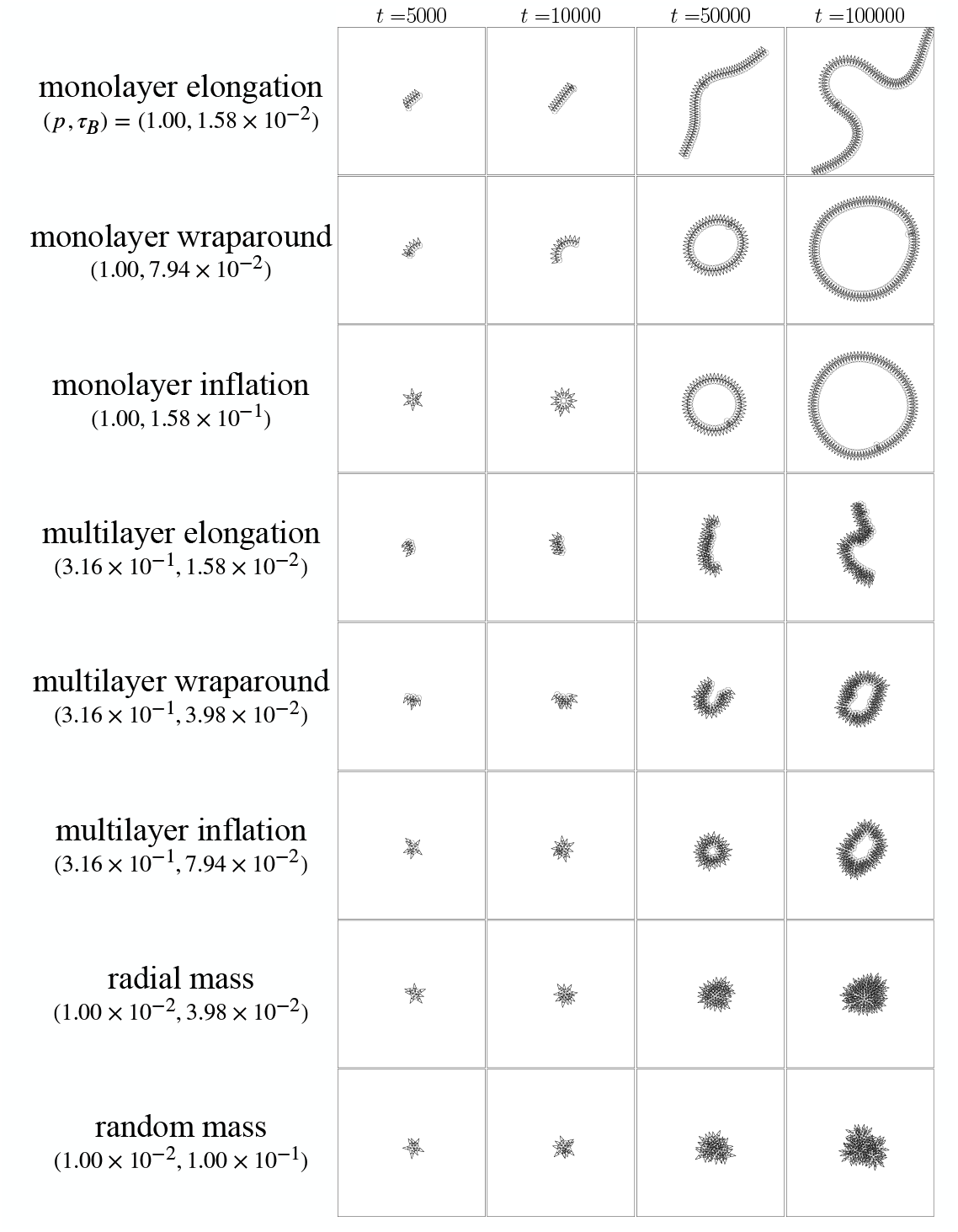
The time series illustrating typical eight behaviors: monolayer elongation, monolayer wraparound, monolayer inflation, multilayer elongation, multilayer wraparound, multilayer inflation, radial mass, and random mass. Each row corresponds to one phase, with the system parameters (*p, τ_B_*) indicated below the corresponding labels. Circles represent the cells, and arrows represent the polarity.

#### Monolayer elongation

Cells align in a row perpendicular to the polarity to form a monolayer. The newly generated cells enter the layer, and the layer extends horizontally. As the line grows longer, limited cell proliferation leads to the “buckling” instability to bend the layer at *t* ≈ 5 × 10^4^. This, however, is beyond our concern.

#### Monolayer wraparound

When *τ*_*B*_ is slightly increased, the cells align in a row as in the previous section, but they extend with a certain curvature and finally stick together at the ends to form a circle.

#### Monolayer inflation

When *τ*_*B*_ is further increased, the cells are still aligned in a row, but they inflate from a lumped state to form a circle.

#### Multilayer elongation

When *p* is smaller than that of the monolayer cases, cells form multiple layers aligned with the polarity and extend horizontally as a whole.

#### Multilayer wraparound

Even in the multilayer region, wraparound was observed.

#### Multilayer inflation

When *τ*_*B*_ was further increased, inflation of a multilayer circle was observed.

### Radial mass

When *p* is much smaller, a mass aligned with the polarity radially is formed without a hole in the center.

### Random mass

When *τ*_*B*_ is large in the multilayer region, a mass with misaligned polarity is formed.

### 3.2 Classification of the behaviors and the phase diagram of the model

The morphologies observed in our model can be classified into eight distinct types. They are distinguished by four main criteria: (i) the degree of polarity alignment, quantified by the time-averaged polarity correlation; (ii) the presence or absence of a central hole and the manner in which it forms; (iii) the magnitude of the mean polarity vector, which reflects the extent of global polarity; and (iv) the number of layers in the resulting structure. A schematic overview of this classification is shown in Fig. 5, along with representative snapshots of each type.

**Fig 5.**
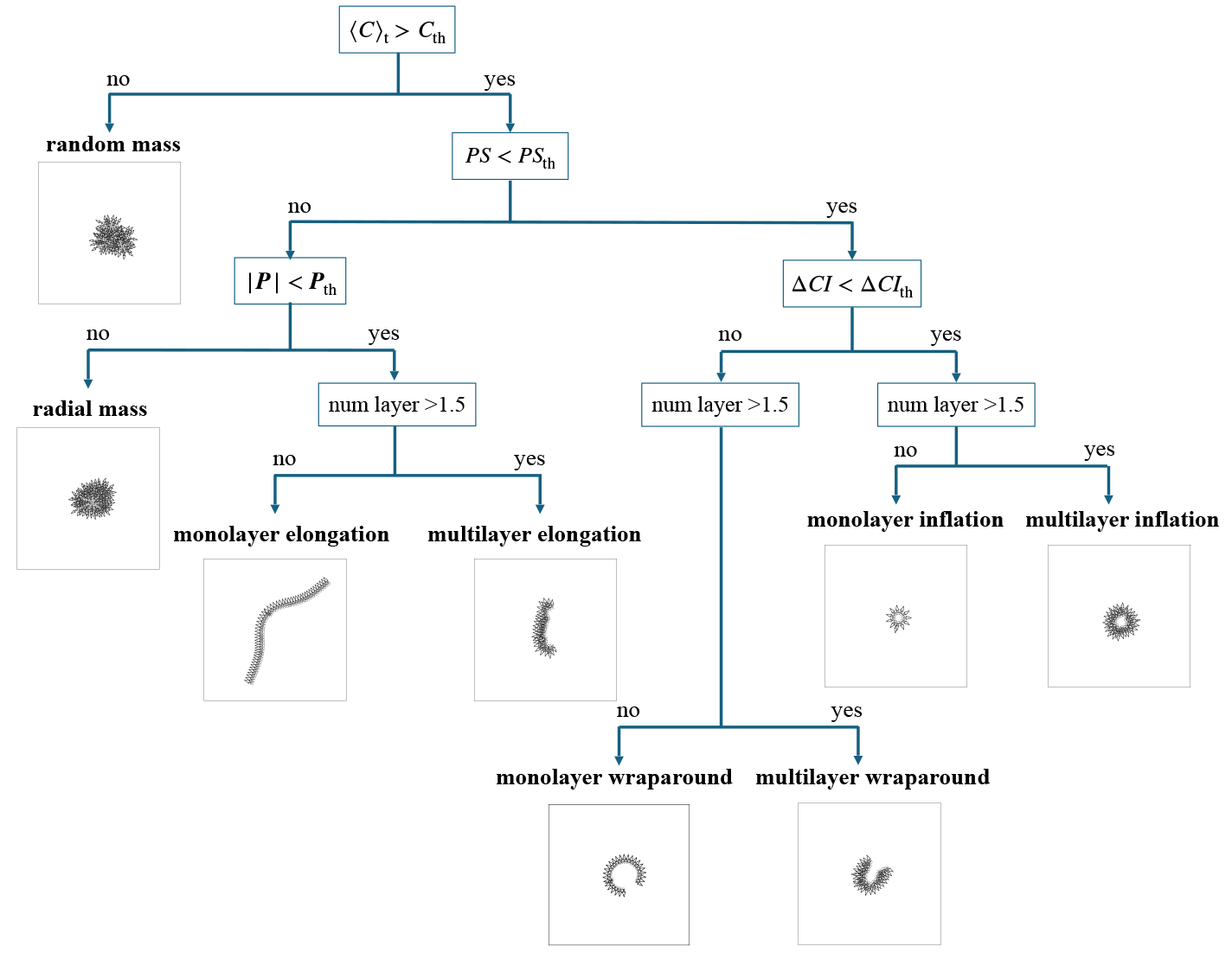
Flowchart of the classification of morphologies observed in this model. Representative behaviors of each morphology are shown under the names of the behaviors.

To make these criteria quantitative, we introduce the following quantities. The polarity correlation ( ⟨*C _t_*⟩) measures how strongly neighboring cells align their polarities over time. The persistence *PS*, calculated using the method of persistent homology, characterizes whether a stable hole is maintained within the structure. The mean polarity magnitude ( |***P***|) indicates the degree of overall polarity order across the system. Finally, the change in circularity index at the moment of hole formation (Δ*CI*) distinguishes whether the hole grows by inflation or by wraparound closure. Details of the definitions and computational procedures of these quantities are provided in the Supporting Text S1.

Based on these quantities, the classification proceeds as summarized in Fig. 5. Configurations with low polarity correlation are categorized as “random mass.” Among the remaining cases, the presence or absence of a hole is then evaluated by *PS*. Structures without a hole are further divided by the magnitude of the mean polarity vector: those with low |***P***|are termed “radial mass,” and those with sufficiently high |***P***|are classified into elongation types. The elongation types are further separated into “monolayer” and “multilayer” depending on the number of layers (with the boundary at 1.5). For hole-containing structures, the distinction between “inflation” and “wraparound” is based on Δ*CI*, and each of these is also subclassified into monolayer and multilayer cases depending on the layer number.

The results presented below were analyzed using the following threshold settings: *C*_th_ = 0.8, *PS*_th_ = 3, *P*_th_ = 0.3, and Δ*CI*_th_ = 1. These values were chosen to effectively separate the qualitatively different behaviors, and small changes in the thresholds do not significantly affect the classification.

Summarizing these behaviors, we obtain the phase diagram shown in Fig. 6. Each grid has 50 samples. We define the representative behavior of the grid as the one that appears most frequently in the 50 samples.

**Fig 6.**
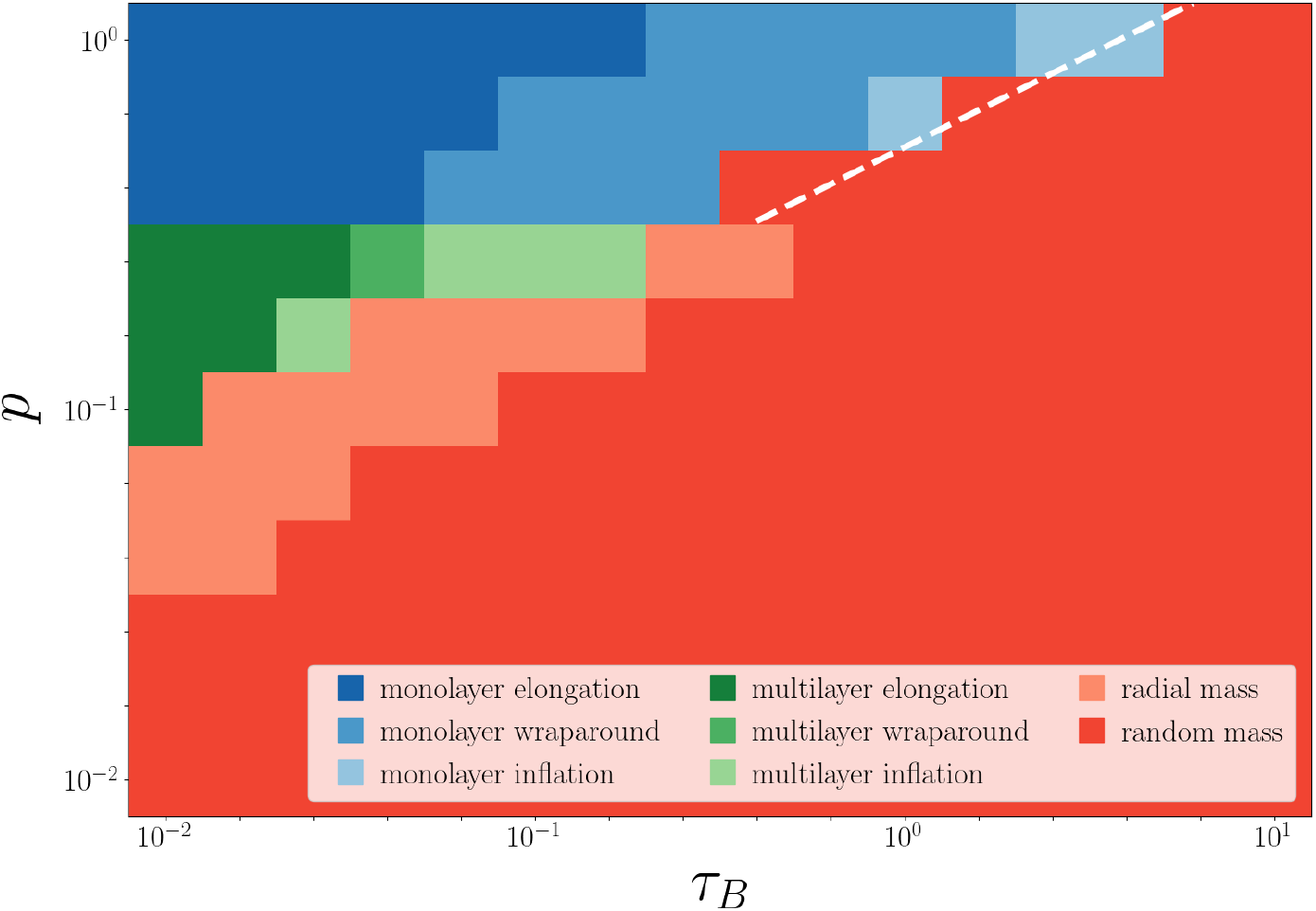
The phase diagram based on *p*, magnitude of polarity, and *τ*_*B*_, the time scale of angle change. Each color represents the behavior shown in the legend. For each parameter set, 50 independent simulation runs were performed, and the most frequently observed behavior is shown by the color of the cell. The white line is obtained from the theoretical estimate of Eq. (18).

First, monolayer and multilayer phases are separated by the magnitude of *p*, of *p*≈ 0.3. Within the multilayer phase at *p* ≳ 0.3, the number of layers increases as *p* increases (Fig. 7). In addition, wraparound and inflation are distinguished by the magnitude of *τ*_*B*_, although the boundary between them depends on *p*. The random mass phase appears when *p* is small or *τ*_*B*_ is excessively large. Note that wraparound is not observed when *τ*_*B*_ is small in the monolayer phase simply because the curvature is too small for the ends of the layer to connect before buckling occurs.

**Fig 7.**
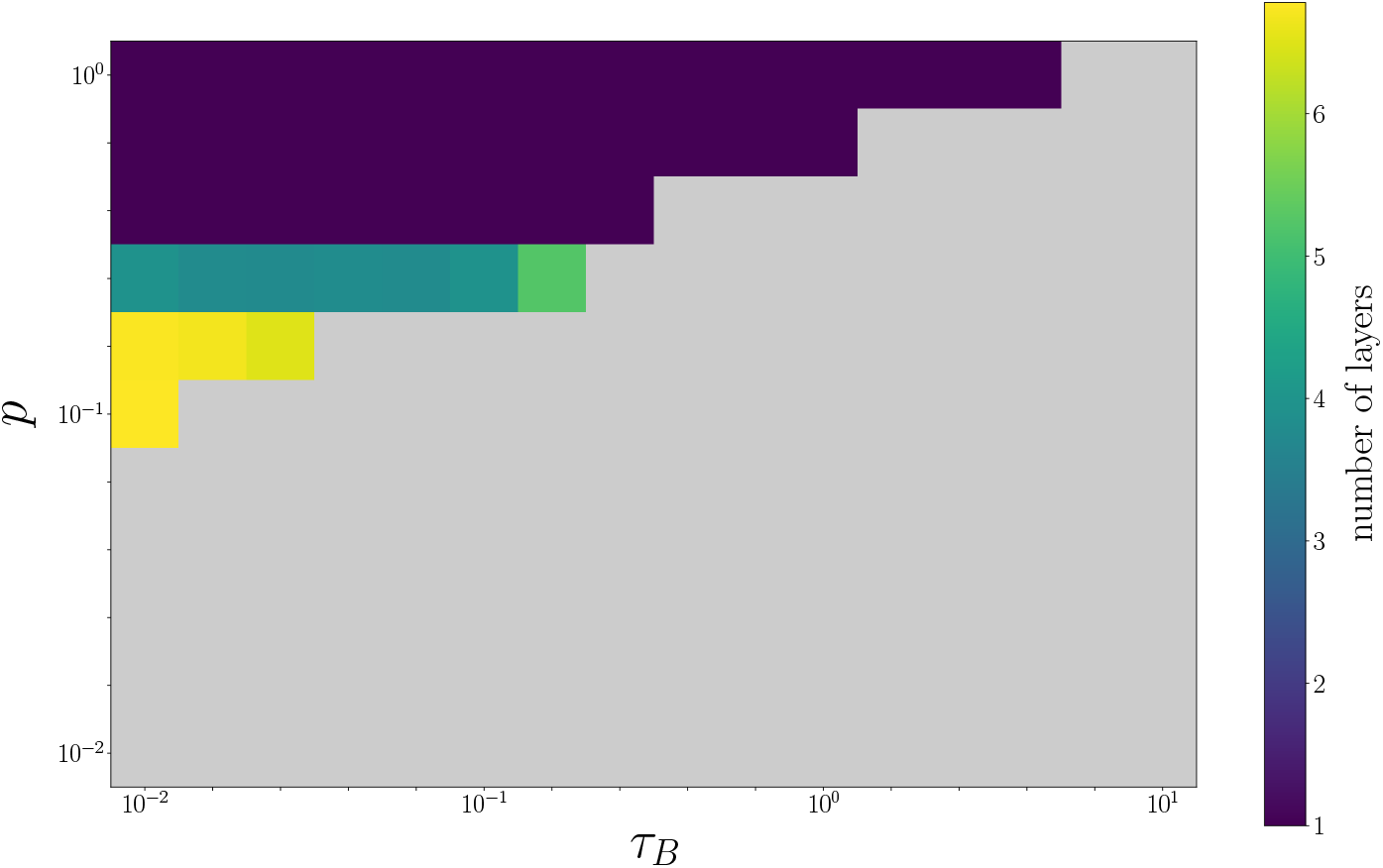
Dependence of the number of layers on *p* and *τ*_*B*_. The color of each grid represents the average number of layers observed, while the light gray regions correspond to the random-mass or radial-mass phases and are excluded from consideration.

From this phase diagram, the parameter *p*, which controls the strength of cell-cell adhesion, determines whether cells form a monolayer or a multilayer. The second key parameter, *τ*_*B*_, controls how strongly neighboring cells tend to orient their polarity in opposite directions. As *τ*_*B*_ increases, the macroscopic dynamics undergo a transition from wraparound to inflation. Thus, wraparound phase results when *τ*_*B*_ is small, while sufficiently large *τ*_*B*_ values lead to inflation. If *τ*_*B*_ is too large, polarity is not oriented at all and the system collapses into the random mass phase.

The phase boundaries depend not only on *τ*_*B*_ but also on *p*. Moreover, inflation is also observed in the multilayer phase, but the required *τ*_*B*_ is much smaller than in the monolayer case. The random mass phase, in turn, appears either when adhesion is weak (*p* small) or when *τ*_*B*_ is excessively large.

To summarize, the collective behavior of cells with polarity-dependent adhesion falls into one of these eight categories depending on *p* and *τ*_*B*_. This result suggests that by considering only the magnitude of polarity and its repulsive effect on adhesive surfaces, basic morphogenetic patterns emerge, and no other patterns exist.

### 3.3 Analysis of phase boundaries

How do these differences in phases appear? To understand the reasons behind the phase diagram, we perform a bifurcation analysis by reducing the system to a few cells to study the transitions between mono-multi layers and wraparound-inflation in the monolayer phase.

#### Analysis of bifurcation between monolayer and multilayer

In this subsection, we focus on the change between monolayer and multilayer in the phase diagram to understand it in terms of bifurcation.

Here, we consider a system consisting of three cells with their polarities in the same direction. In this analysis, we set a line connecting the centers of cells 1 and 2 as the *x*-axis, and the midpoint of these two cells as the origin (Fig. 8a). We examine if the cell 3 moves upward leading to a monolayer structure or not. Specifically, we analyze whether the third cell goes upward and is aligned with the first and second cells to form a one-dimensional chain in the stable state. From the symmetry of the interaction received from cells 1 and 2, the cell 3 does not move in the x-direction whereas it can move along the *y*-axis perpendicular to the *x*-axis. Then, the dependence of the cell configuration on the parameter *p* is studied in terms of bifurcation between mono- and multilayer.

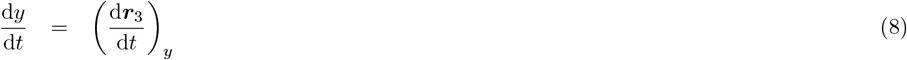

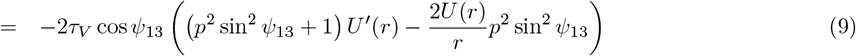

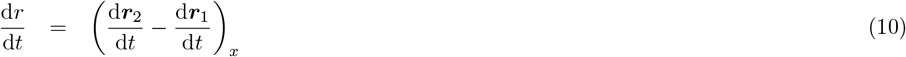

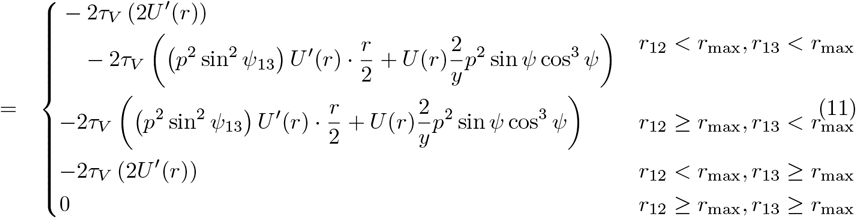

**Fig 8.**
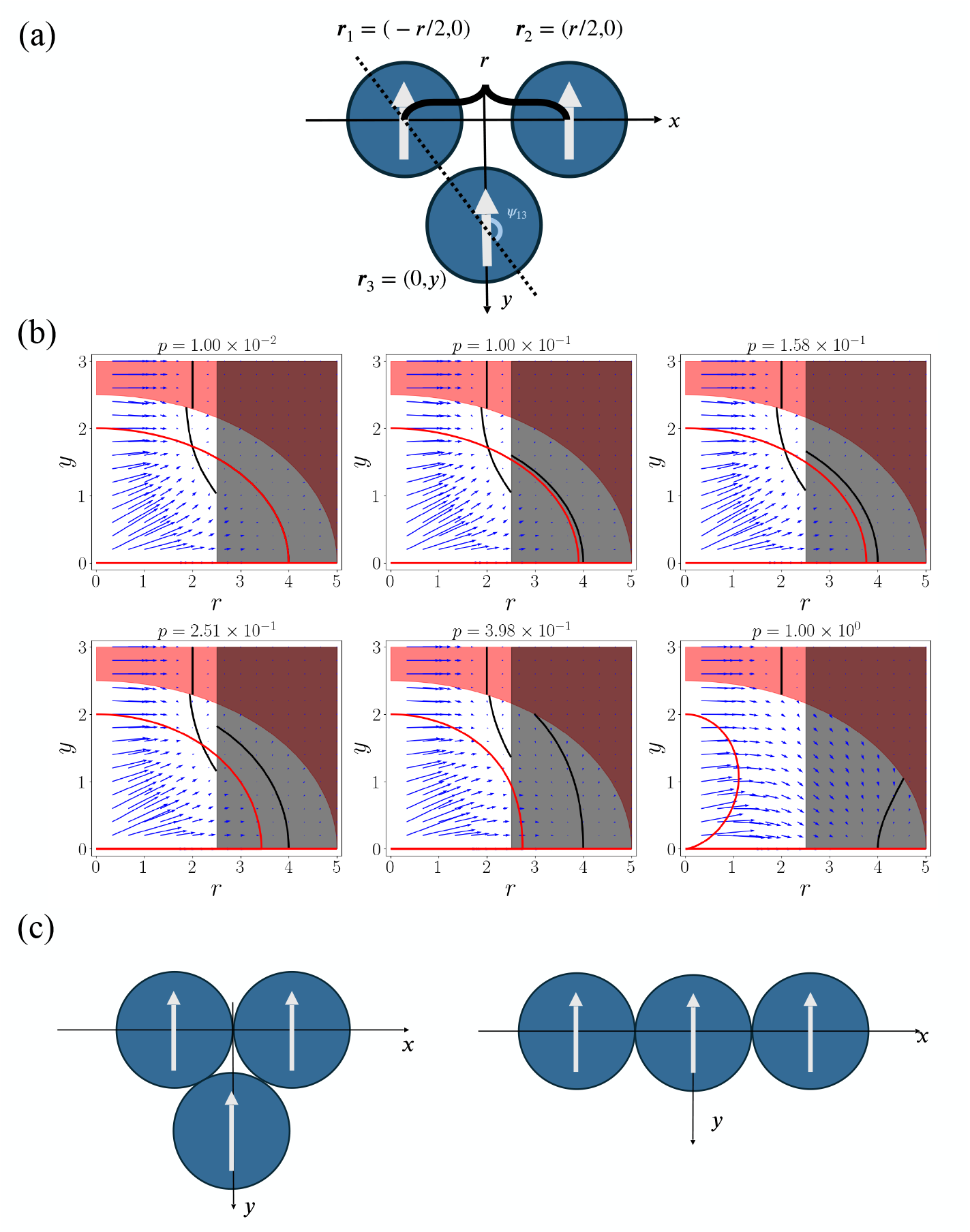
Three-cell bifurcation analysis of the monolayer–multilayer transition. (a) In this analysis, we consider a system of three cells interacting with each other, all having the same polarity direction. When cells 1, 2, and 3 are located as shown in the figure, we investigate how the initial positions determined by *r* and *y* influence the final configuration of the three cells from a dynamics perspective. (b) Differences in the dynamics of *r* and *y* and the resulting fixed points for each *p*. The blue arrows represent the velocity field, while the black and red lines represent the nullclines of *r* and *y*, respectively. The black and red shaded areas correspond to *r*_12_ *> r*_max_ and *r*_13_ *>r*_max_, respectively. (c) Left: Cell configuration at the fixed point 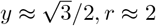. Right: Cell configuration at the fixed point *y* = 0, *r* = 4.

Here, sin 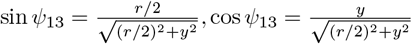.

To analyze the system of (*r, y*), we consider two nullclines d*y/*d*t* = 0 and d*r/*d*t* = 0. Although analytic representation is difficult to obtain, they are drawn numerically as in Fig. 8b. The result is shown in Fig. 8b with the velocity field.

When the value of *p* is large (*p* ≥0.5), this system has only one stable fixed point with *y* = 0 and *r* = 4. This means that all three cells are aligned within the same axis (Fig. 8c). As *p* decreases, there is a saddle-node bifurcation at *p* ≈ 0.3 so that this system with smaller *p* has two stable fixed points with (*y* = 0, *r* = 4) and 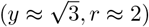. The latter fixed point means that the third cell is located below the first two cells (Fig. 8c). The existence of this fixed point allows the system to form a multilayer. This bifurcation point at *p* ≈ 0.3 roughly agrees with the boundary in the phase diagram in Figure 6. Note that the fixed point with *y* = 0 and *r* = 4 is still stable in this region.

Since the dynamics of the polarity *p* is neglected, this analysis does not strictly explain the entire boundary between monolayer and multilayer shown in Fig. 7. In this derivation, we assumed that the polarity of the three cells is aligned, and the dynamics of ***p***(*θ*) in the actual model is not considered. If the dynamics of ***p***(*θ*) in the actual model is considerably different from this assumption, the analysis in this section is not valid. For example, in the monolayer-inflation phase where *τ*_*B*_ is large, the polarities of the three cells are radially aligned in the steady state of the three cells. This is beyond the scope of the discussion in this section. However, in the region where *τ*_*B*_ ≤ 0.2, this assumption on aligned polarity is valid to some extent, according to the numerical observation. Therefore, at least for the boundary between monolayer and multilayer regions, the present estimate of phase boundary is valid.

### Analysis of the boundary between monolayer wraparound and inflation

Next, we analyze the phase boundary between wraparound and inflation, in terms of cell-cell interactions. We focus on the change between wraparound and inflation in monolayer case [29]. Here, we consider a three-cell system, and examine whether the cells at both ends interact or not. This is because when the cells at both ends do not interact, a system with more than three cells can be decomposed into consecutive adjacent two-cell systems, leading to angles between them needed for a wraparound structure. Now, assume that the ends do not interact with each other. In this case, the three-body system can be represented by two adjacent two-body systems (Fig. 9). The fixed points of a two-body system can be derived as follows.

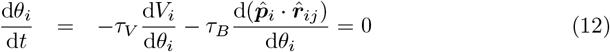

**Fig 9.**
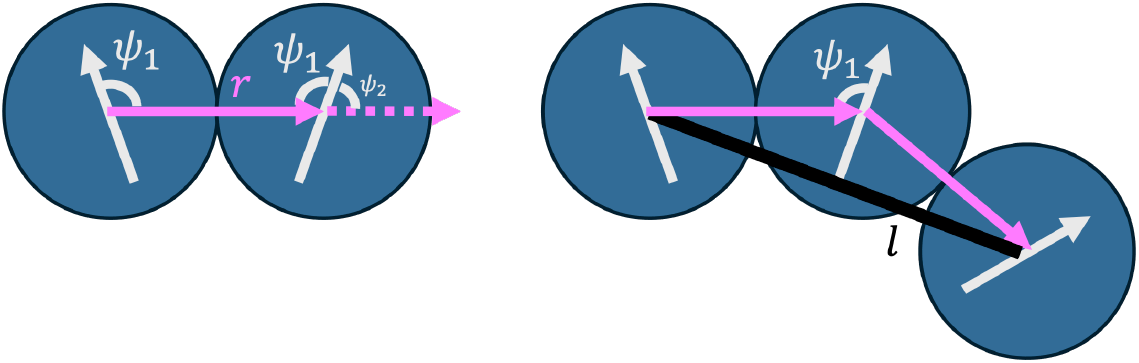
Two-body reduction of a three-cell system for the wraparound–inflation boundary. In this analysis, we focus on whether the two end cells interact with each other when three cells are aligned while maintaining the polarity angles of the two-cell system. The angles of the polarity vectors with respect to *r* in the two-cell system are denoted as *ψ*_1_ and *ψ*_2_ in the figure. Due to symmetry, *ψ*_2_ = *π* −*ψ*_1_. The distance between the two end cells is represented by the black line, and its length is denoted as *l*.

Since the two-body system is described by two degrees of freedom *θ*_1_ and *θ*_2_, by solving Eq. (12) for *θ*_1_ and *θ*_2_, we get

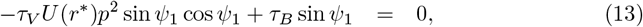

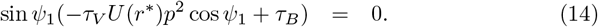

Therefore, Eq. (12) holds when *ψ*_1_ satisfies either of the following conditions.

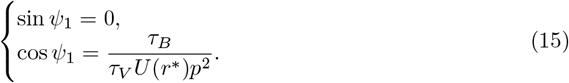

When these two-body systems are connected, the distance between the ends can be calculated. Therefore, assuming the stable solution [30], the condition that the ends of the cells do not interact with each other is written as

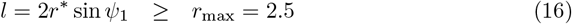

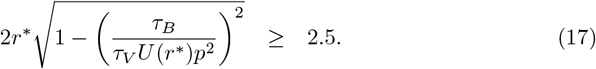

Substituting the values used in the simulation *r*^∗^ = 2, *U* (*r*^∗^) 0.5, and *τ*_*V*_ = 10 into the equations, we get:

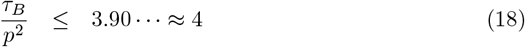

Within the range of the above inequality, a wraparound is expected to occur. This can explain the trend of the boundary between wraparound and inflation in the phase diagram (Fig. 6).

### 3.4 3D version of the model

So far, we have presented the results for the two-dimensional case. However, the model can be straightforwardly extended to a three-dimensional case. In the three-dimensional case, we also confirmed the same types of morphogenesis as those observed in the two-dimensional model, except for the random mass phase. These are shown in Supporting Figures (Figs. S12-S18). In three dimensions, there is one more degree of freedom regarding the direction of polarity compared to two dimensions, which is expected to amplify the polarity alignment so that random mass phase will be unlikely to exist. Except for this, we have the seven phases as observed in the two-dimensional case, with similar dependence on *p* and *τ*_*B*_.

### 3.5 Possible extension of the model to correspond to actual morphogenesis

In the previous sections, we demonstrated that basic types of cell configuration (Fig. 6) are represented as distinct “phases” with the parameters for adhesion and polarity. In the real developmental process, these parameters are not necessarily identical over all cells, but can depend on the position or on their cell types. In this section, we show that this model can express more complex morphologies by combining the results of each phase. We show that the early embryos of fish and mammals are reproduced by simply introducing the spatial gradient of *τ*_*B*_ or time evolution of *p*.

#### Gradient of *τ*_*B*_ along *y*-axis: as a possible model for fish morphogenesis

In the early embryos of fish, cells generally form a multilayer wraparound. However, within this process, cells at the leading edge are drawn inward as they are engulfed [1]. This behavior can be reproduced by assuming that each cell has a different *τ*_*B*_ value depending on its position. As shown in Fig. 10, by setting a gradient in *τ*_*B*_ along the *y*-axis, the effective curvature radius becomes smaller in regions with high *τ*_*B*_, allowing the leading edge cells to be drawn inward [31]. This corresponds to the characteristic morphogenetic movement in fish epiboly, where blastodermal cells undergo inward migration while simultaneously advancing over and enclosing the yolk.

**Fig 10.**
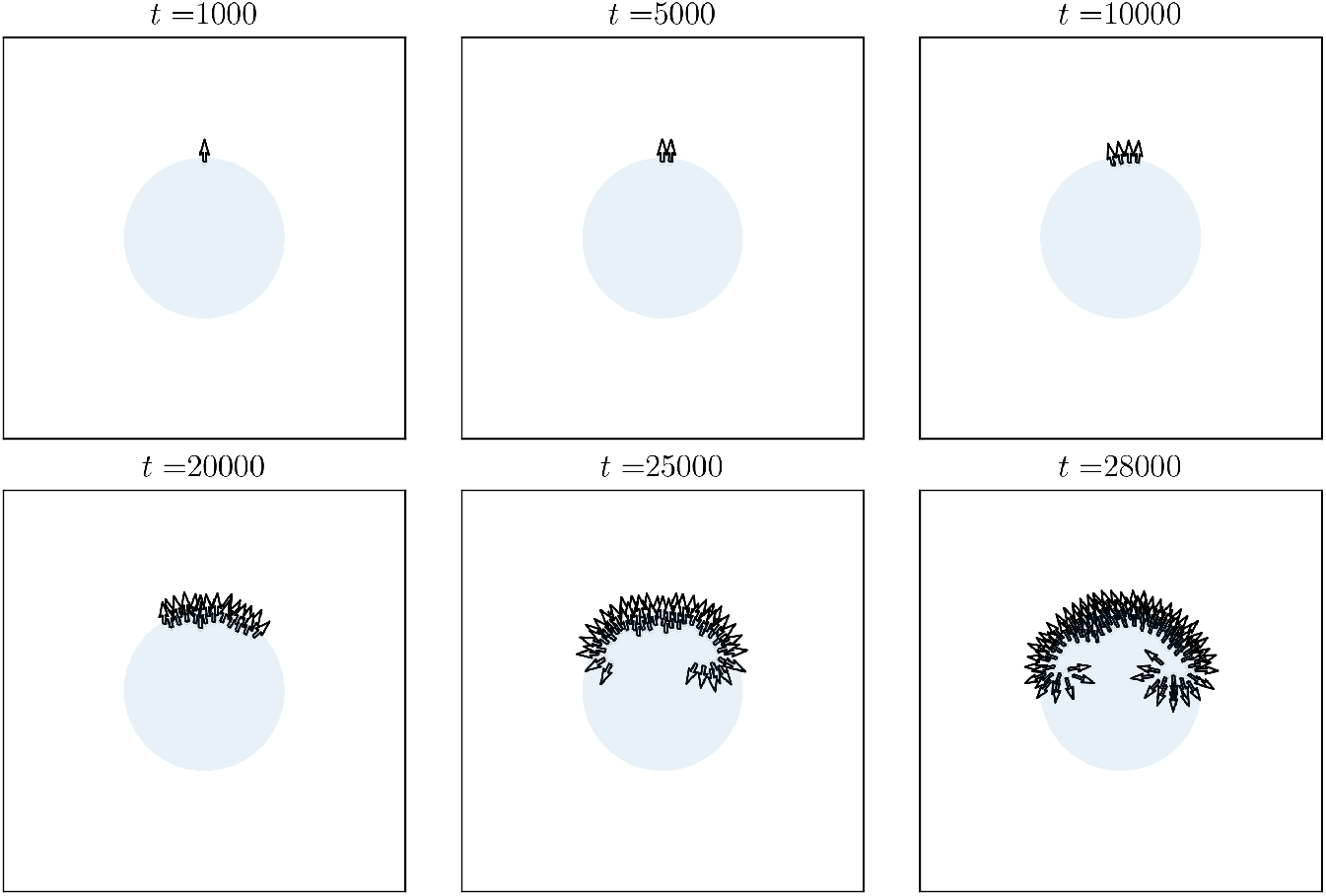
Morphogenesis process with a gradient in *τ*_*B*_. Here, the strength of polarity follows *τ*_*B*_ = 2 − 0.1(*y*_*i*_ − 50).

#### Time-dependent *p* driven by a diffusive signal: as a possible model for mammalian morphogenesis

The early embryos of mammals initially form a solid mass, which then differentiates into an outer monolayer sphere and an inner cell mass. This behavior can be reproduced by allowing *p* to change over time in response to an external signal. As shown in Fig. 11, the outer cells are exposed to the signal and acquire polarity, exhibiting monolayer inflation behavior. Whereas the outer cells acquire polarity, the inner cells cannot receive the signal because the outer cells act as a barrier, so they keep low polarity and stay in the random mass phase. This corresponds to the characteristic morphogenetic event in early mammalian development, where outer cells organize into a single-layered epithelium surrounding an inner cell mass that remains inside.

**Fig 11.**
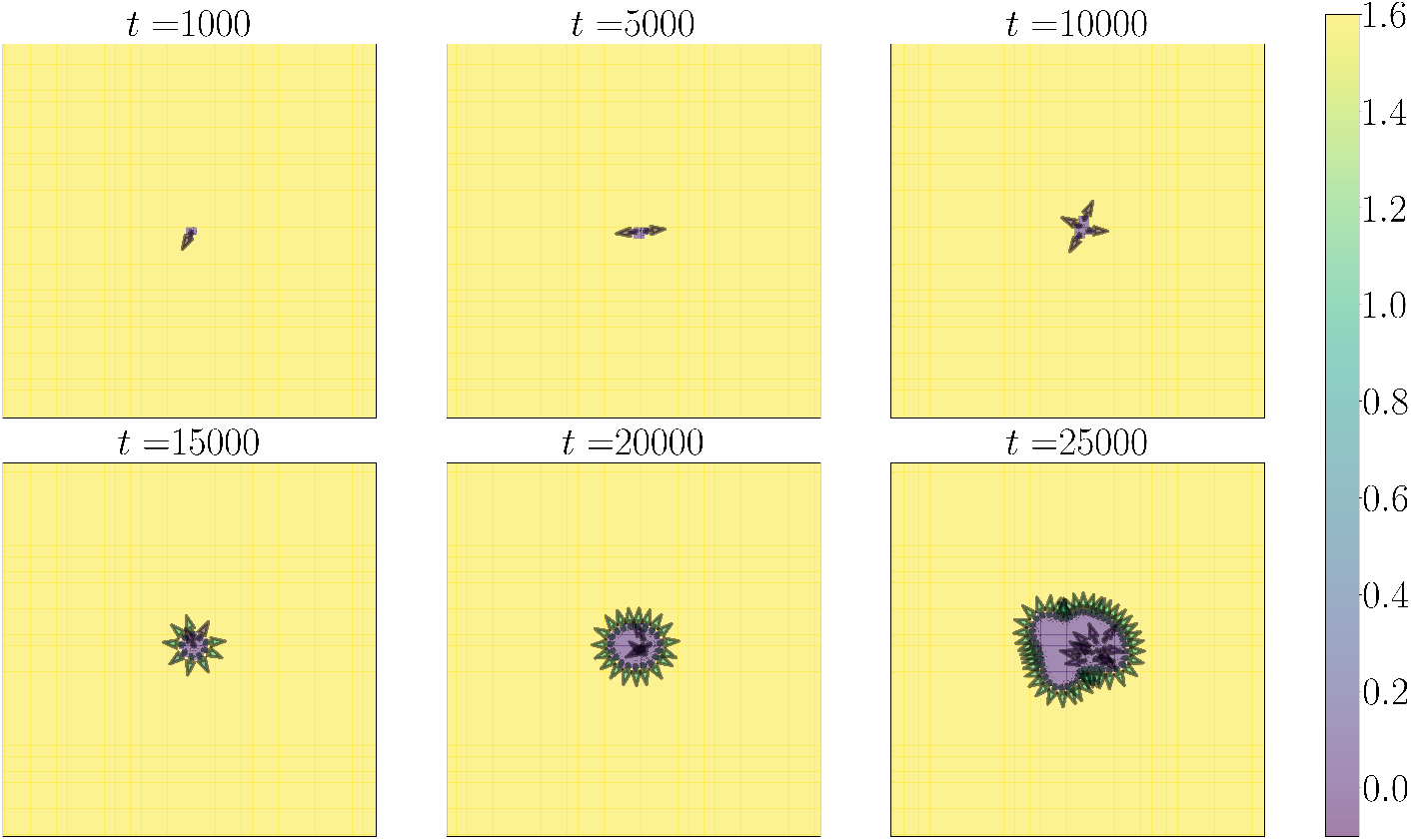
Morphogenesis process when *p* develops over time due to a diffusive signal. The signal does not diffuse within the field occupied by the cells. Cells absorb the signal from adjacent regions and determine their polarity strength while consuming the signal. The color of the field represents the amount of signal, and the color of the arrows represents the strength of each cell’s polarity. For details of the model, refer to Supporting Text S2.

## 4 Discussion

In this study, we used a mathematical model to investigate the role of polarity in the formation of cellular layers, masses, and cavities. Despite its simplicity, our model incorporating only adhesion and polarity is able to replicate the basic types of morphogenetic patterns observed in early embryogenesis. A key feature of the model is that the observed patterns are governed by just two parameters. The first one is *p*, which represents the strength of polarity and thereby controls the extent to which polarity drives structural organization. The second one is *τ*_*B*_, which specifies the time scale of polarity regulation, i.e., how rapidly polarity orientations change; in our analysis, *τ*_*V*_ was fixed and *τ*_*B*_ was varied to assess its effect. By focusing on these two parameters, the diversity of morphogenetic behaviors is generated by the model, as can be systematically understood. Our findings revealed that the number of cellular layers increases with the polarity parameter *p*, transitioning from monolayer to multilayer configurations. Additionally, we identified two distinct mechanisms of cavity formation: wraparound and inflation. The mode of cavity formation is determined by the time scale of polarity regulation (*τ*_*B*_), highlighting the critical role of polarity regulation in cavity development within multicellular systems. Overall, our model provides a theoretical framework for understanding the morphogenesis of multicellular systems and offers insights into how polarity and its regulation contribute to the structural complexity observed in real organisms.

Our model highlights the importance of the time scale of polarity regulation in determining the mode of cavity formation. This effect described by the second term in the equation (7) is based on the assumption that cell adhesion influences the orientation of polarity. In fact, examples of polarity formation observed in various developmental processes [33, 34] and organoids [35] clearly indicate that adhesion exerts some control over polarity, typically by orienting the apical side distinct from adhesion sites. Given these biological observations, we consider our modeling assumption to be qualitatively natural. Fundamental morphogenetic patterns can be understood as generic physical processes [9], in the spirit of universal biology [32].

In this model, qualitative differences in macroscopic morphogenesis can be explained by the combination of only two microscopic parameters: the magnitude of polarity *p* and the time scale of angle change *τ*_*B*_. As shown in Fig. 6, distinct morphogenetic modes, such as multilayer inflation and monolayer wraparound, emerge in adjacent regions of the (*p, τ_B_*) parameter space. This indicates that just a change in these parameters can trigger a drastic transition in the global morphogenetic behavior. Consequently, when interpreting evolutionary changes in developmental processes through this model, similarity in macroscopic morphology may not necessarily correspond to phylogenetic proximity. This result from the model is consistent with observations that amphibians and mammals generally exhibit inflation-like morphogenesis in the early embryogenesis, whereas many fish (including zebrafish) and birds typically show wraparound-like patterns.

Our results provide a new perspective on the role of polarity in morphogenesis, which can be validated and observed through experiments. By effectively altering the parameters *p* and *τ*_*B*_, it may be possible to change the mode of morphogenesis, according to the phase diagram in Fig. 6. For example, cadherins are known to mediate cell-cell adhesion and play a critical role in maintaining tissue integrity and structure [11]. By manipulating the expression levels of cadherins, researchers have observed changes in tissue morphology, supporting the idea that cell adhesion strength can influence macroscopic morphology [12, 36]. Furthermore, when E-cadherin is knocked out, mouse ES cells form a monolayer sphere by wrapping around, corresponding to the monolayer wraparound phase in our study; otherwise, they typically form a monolayer sphere by inflating from within during proliferation, which corresponds to the monolayer inflation phase [37, 38]. This can be interpreted as a transition from the monolayer inflation phase to the monolayer wraparound phase in our study. By manipulating the expression levels or activity of such adhesion molecules, one could potentially influence the polarity dynamics and the resulting morphological outcomes. This suggests that the regulation of *p* and *τ*_*B*_ could serve as a mechanism to control the formation of different tissue structures or the number of layers, providing a valuable tool for both developmental biology and tissue engineering. In future studies, a more extensive exploration of the parameters will be necessary.

Here, we considered a model where the total number of cells increases linearly, but of course this is not always the case in reality. For example, if each cell divides with a constant cell cycle, the number of cells will increase exponentially. In this scenario, the number of new cells added to the system also increases exponentially, meaning that after a certain period, new cells are added before the adhesion dynamics have converged to a stable state. This can lead to phenomena such as buckling later, which differ from the results observed in this study. This type of dynamics that occur when cell proliferation is rapid are beyond the scope of this research. The reason why the number of layers changes in this model is still not yet explained analytically. It seems that, in principle, any number of layers could be stacked if the three-cell system possesses a stable fixed point that allows a third cell to pile on top (Fig. 8c). However, our simulations show that the actual number of layers is limited and is determined at the macroscopic level rather than solely by local three-cell interactions. As a hypothesis, the number of layers may depend on the relative sizes of the basins of attraction corresponding to different stable configurations in the three-cell system. In other words, the probability that a third cell stabilizes on top of or within the existing layer may determine the macroscopic number of layers. Clarifying this relationship remains a topic for future research. Additionally, we were unable to analytically understand the boundary between wraparound and inflation in the multilayer case, which also depends on the value of *p*. This result remains a subject for future theoretical research as well.

The potential applications of our model are not limited to the morphogenesis of current organisms. The insights gained from our study can be useful for designing artificial multicellular systems, such as organoids, artificial meat, and regenerative medicine. Organoids, which are three-dimensional structures derived from stem cells that mimic the architecture and function of real organs, have become valuable tools for studying development and disease [39]. By applying our model to organoid systems, one can identify optimal culture conditions and manipulation of cell polarity to achieve desired tissue structures.

## Code Availability

All codes used for analysis and visualization are available at https://github.com/Nakayoshi98/Adhesion-and-polarity-driven-morphogenesis

## Acknowledgments

This work was supported by RIKEN Junior Research Associate Program (to YTN), the Japan Society for Promotion of Science (JSPS) KAKENHI (22K21344 and 24H01798 to CF), and the Novo Nordisk Foundation (Grant No.NNF21OC0065542 to KK). We thank K. Sneppen, A. Trusina and M. Kitazawa for stimulating discussions.

## Notes

### Competing Interest Statement

The authors have declared no competing interest.

## References

1. Wolpert L, Tickle C, Arias AM. Principles of development. Oxford University Press, USA; 2019.

2. Kimmel CB, Ballard WW, Kimmel SR, Ullmann B, Schilling TF. Stages of embryonic development of the zebrafish. Developmental Dynamics. 1995;203(3):253–310. doi:10.1002/aja.1002030302.

3. Bruce AEE. Zebrafish epiboly: Spreading thin over the yolk. Developmental Dynamics. 2016;245(3):244–258. doi:10.1002/dvdy.24353.

4. Slack JMW. In vitro development of isolated ectoderm from axolotl gastrulae. Development. 1984;80(1):321–330. doi:10.1242/dev.80.1.321.

5. Johnson MH, McConnell JML. Lineage allocation and cell polarity during mouse embryogenesis. Seminars in Cell & Developmental Biology. 2004;15(5):583–597. doi:10.1016/j.semcdb.2004.04.002.

6. Giudice G. Developmental biology of the sea urchin embryo. Elsevier; 2012.

7. Fairclough SR, Dayel MJ, King N. Multicellular development in a choanoflagellate. Current Biology. 2010;20(20):R875–R876. doi:10.1016/j.cub.2010.09.014.

8. Brunet T, King N. The Origin of Animal Multicellularity and Cell Differentiation. Developmental Cell. 2017;43(2):124–140.

9. Forgacs G, Newman SA. Biological physics of the developing embryo. Cambridge University Press; 2005.

10. Newman SA, Bhat R. Dynamical patterning modules: a “pattern language” for development and evolution of multicellular form. Int J Dev Biol. 2009;53(5-6):693–705. doi:10.1387/ijdb.072481sn.

11. Takeichi M. Cadherin Cell Adhesion Receptors as a Morphogenetic Regulator. Science. 1991;251(5000):1451–1455. doi:10.1126/science.2006419.

12. Gumbiner BM. Regulation of cadherin-mediated adhesion in morphogenesis. Nature Reviews Molecular Cell Biology. 2005;6(8):622–634. doi:10.1038/nrm1699.

13. Nose A, Nagafuchi A, Takeichi M. Expressed recombinant cadherins mediate cell sorting in model systems. Cell. 1988;54(7):993–1001. doi:10.1016/0092-8674(88)90114-6.

14. Hatta K, Takagi S, Fujisawa H, Takeichi M. Spatial and temporal expression pattern of N-cadherin cell adhesion molecules correlated with morphogenetic processes of chicken embryos. Developmental Biology. 1987;120(1):215–227. doi:10.1016/0012-1606(87)90119-9.

15. Mogilner A, Allard J, Wollman R. Cell polarity: quantitative modeling as a tool in cell biology. Science. 2012;336(6078):175–179. doi:10.1126/science.1216380.

16. Buckley CE, St Johnston D. Apical–basal polarity and the control of epithelial form and function. Nature Reviews Molecular Cell Biology. 2022;23(8):559–577. doi:10.1038/s41580-022-00465-y.

17. Goldstein B, Macara IG. The PAR Proteins: Fundamental Players in Animal Cell Polarization. Developmental Cell. 2007;13(5):609–622. doi:10.1016/j.devcel.2007.10.007.

18. Rodriguez-Boulan E, Macara IG. Organization and execution of the epithelial polarity programme. Nature Reviews Molecular Cell Biology. 2014;15(4):225–242. doi:10.1038/nrm3775.

19. Motegi F, Plachta N, Viasnoff V. Novel approaches to link apicobasal polarity to cell fate specification. Current Opinion in Cell Biology. 2020;62:78–85. doi:10.1016/j.ceb.2019.09.003.

20. Graner F, Glazier JA. Simulation of biological cell sorting using a two-dimensional extended Potts model. Phys Rev Lett. 1992;69(13):2013–2016. doi:10.1103/PhysRevLett.69.2013.

21. Glazier JA, Graner F. Simulation of the differential adhesion driven rearrangement of biological cells. Phys Rev E. 1993;47(3):2128–2154. doi:10.1103/PhysRevE.47.2128.

22. Belmonte JM, Clendenon SG, Oliveira GM, Swat MH, Greene EV, Jeyaraman S, et al. Virtual-tissue computer simulations define the roles of cell adhesion and proliferation in the onset of kidney cystic disease. Molecular Biology of the Cell. 2016;27(22):3673–3685. doi:10.1091/mbc.E16-01-0059.

23. Honda H. Description of cellular patterns by Dirichlet domains: The two-dimensional case. J Theor Biol. 1978;72(3):523–543. doi:10.1016/0022-5193(78)90315-6.

24. Farhadifar R, Röper JC, Aigouy B, Eaton S, Jülicher F. The influence of cell mechanics, cell-cell interactions, and proliferation on epithelial packing. Curr Biol. 2007;17(24):2095–2104. doi:10.1016/j.cub.2007.11.049.

25. Fletcher AG, Osterfield M, Baker RE, Shvartsman SY. Vertex models of epithelial morphogenesis. Biophys J. 2014;106(11):2291–2304. doi:10.1016/j.bpj.2013.11.4498.

26. Okuda S, Inoue Y, Adachi T. Three-dimensional vertex model for simulating multicellular morphogenesis. Biophysics and Physicobiology. 2015;12:13–20. doi:10.2142/biophysico.12.013.

27. Nissen SB, Rønhild S, Trusina A, Sneppen K. Theoretical tool bridging cell polarities with development of robust morphologies. eLife. 2018;7:e38407. doi:10.7554/eLife.38407.

28. Hagolani PF, Zimm R, Marin-Riera M, Salazar-Ciudad I. Cell signaling stabilizes morphogenesis against noise. Development. 2019;146(20):dev179309. doi:10.1242/dev.179309.

29. Similar analysis would be possible for the multilayer case, but it is difficult to get analytical results.

30. If ψ_1_ that satisfies the second equation for cos ψ_1_ in Eq. (15) exists, that solution is stable.

31. In this model, boundary conditions are set to be circle (gray area in Fig. 10). This is because, in two dimensions, the shape of the multilayer wraparound does not robustly form a perfect circle. If the shape does not form a perfect circle, the effect of the gradient in τ_B_ can differ at both ends, causing inconsistencies. However, in the three-dimensional model, the multilayer wraparound often forms a more perfect sphere. Therefore, we consider this change in boundary conditions to be addressing a two-dimensional specific issue and not problematic.

32. Kaneko K. Universal Biology: The Physics of Life through the Macro-Micro Consistency Principle. Cambridge University Press; 2025.

33. Alford LM, Ng MM, Burgess DR. Cell polarity emerges at first cleavage in sea urchin embryos. Developmental Biology. 2009;330(1):12–20. doi:10.1016/j.ydbio.2009.02.039.

34. Maître JL. Mechanics of blastocyst morphogenesis. Biology of the Cell. 2017;109(9):323–338. doi:10.1111/boc.201700029.

35. Wang H, Lacoche S, Huang L, Xue B, Muthuswamy SK. Rotational motion during three-dimensional morphogenesis of mammary epithelial acini relates to laminin matrix assembly. Proceedings of the National Academy of Sciences. 2013;110(1):163–168. doi:10.1073/pnas.1201141110.

36. Bao M, Cornwall-Scoones J, Sanchez-Vasquez E, Cox AL, Chen DY, De Jonghe J, et al. Stem cell-derived synthetic embryos self-assemble by exploiting cadherin codes and cortical tension. Nature Cell Biology. 2022;24(9):1341–1349. doi:10.1038/s41556-022-00984-y.

37. Liang X, Weberling A, Hii CY, Zernicka-Goetz M, Buckley CE. E-cadherin mediates apical membrane initiation site localisation during de novo polarisation of epithelial cavities. The EMBO Journal. 2022;41(24):e111021. doi:10.15252/embj.2022111021.

38. Herranz G, Martín-Belmonte F. Cadherin-mediated adhesion takes control. The EMBO Journal. 2022;41(24):e112662. doi:10.15252/embj.2022112662.

39. Hofer M, Lutolf MP. Engineering organoids. Nature Reviews Materials. 2021;6(5):402–420. doi:10.1038/s41578-021-00279-y.

40. Post MJ. Cultured meat from stem cells: Challenges and prospects. Meat Science. 2012;92(3):297–301. doi:10.1016/j.meatsci.2012.04.008.

41. Atala A. Tissue Engineering and Regenerative Medicine: Concepts for Clinical Application. Rejuvenation Research. 2004;7(1):15–31. doi:10.1089/154916804323105053.

