## Supporting Information for "Adhesion and polarity-driven morphogenesis: Mechanisms and constraints in tissue formation"

### S1 Text: Morphological features and their dynamics

To quantitatively characterize the behaviors observed in the model, we introduced order parameters in the main text. Below, we describe them in detail, as follows:

1. Polarity correlation: Whether the polarities are aligned.
2. Presence of a hole: Whether there is a hole in the center.
3. Formation process of the hole: How the hole was formed.
4. Number of layers: The number of layers.
5. Mean field of polarity: The mean field of polarity.

#### 1 Polarity correlation

First, to determine whether the polarities are aligned, we use polarity correlation, which is defined below.

$$C = \frac{1}{N} \sum_i \hat{\boldsymbol{p}}_i \cdot \left( \frac{1}{|N_i|} \sum_{j \in N_i} \hat{\boldsymbol{p}}_j \right), \tag{1}$$

where $N$ is the total number of cells, and $N_i$ is a set of cells $j(\neq i)$ such that $r_{ij} < r_{\max}$. Again, $\hat{\boldsymbol{p}}_i$ is the normalized polarity vector of cell $i$.

With this definition, it can be seen in the figures that when cells are orderly aligned in the same direction with their neighbors, the correlation is high (Fig. S1), whereas when they are randomly oriented, the correlation is nearly zero (Fig. S2). We classify the behavior as “random mass”, if the time-averaged polarity correlation does not exceed a given threshold (here 0.8) (the following results do not significantly depend on this choice of value, though).

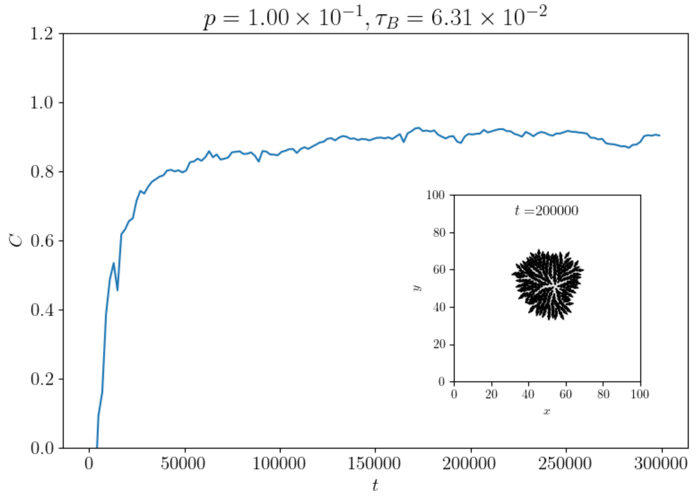


**Fig S1.** Time series of polarity correlation in a single simulation with $p = 1.00 \times 10^{-1}$ and $\tau_B = 6.31 \times 10^{-2}$. The inset shows the arrangement of cells at $t = 200000$, with the polarity of each cell represented by arrows.

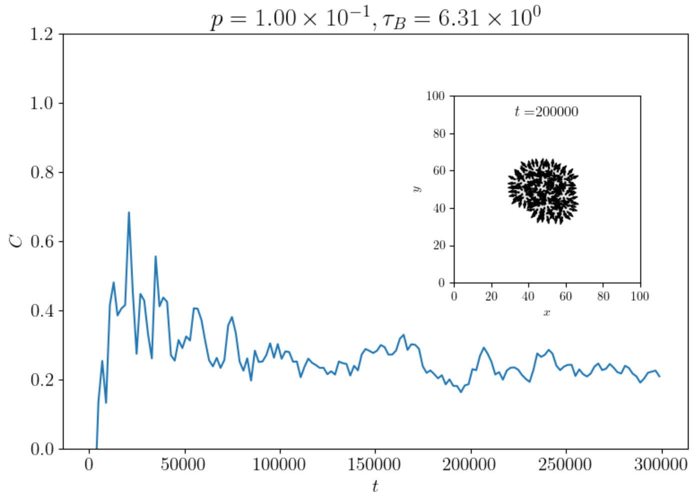


**Fig S2.** Time series of polarity correlation in a single simulation with $p = 1.00 \times 10^{-1}$ and $\tau_B = 6.31 \times 10^{0}$.

### 2 Presence of a hole

We use persistent homology [1] to determine the presence of a hole.

Persistent homology can be intuitively understood by considering a set of data points in space, where we gradually increase the radius ($R$ in Fig. S3) of circles centered at each point. Initially, when the radius is small, each point is isolated, and no significant topological features are present. As the radius increases, the circles begin to overlap, and connected components start to form.

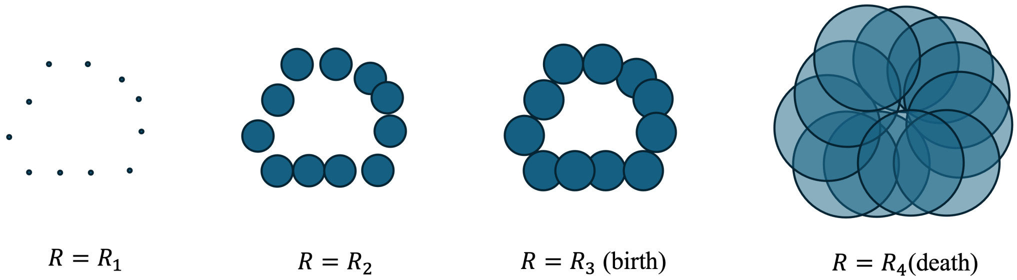


**Fig S3.** Schematic description of persistent homology. As the radius of the circle increases, connected components begin to form.

As we continue to increase the radius, some of these connected components merge, and one-dimensional holes begin to appear. The key idea is to track the birth and death of these holes as the radius changes. The "birth" of a hole is the radius at which it first appears ($R_3$ in Fig. S3), and its "death" is the radius at which it merges with another hole or disappears ($R_4$ in Fig. S3). The "persistence" of a hole is the difference between its birth and death radii ($R_4 - R_3$), which is intuitively regarded as the size of the hole.

Here, we consider only those features with a birth radius smaller than $R_{\text{cell}} = 1.25$, since the typical radius of a cell is 1 [2]. The largest persistence of the holes that satisfy this condition is used as the size of the largest hole in the system. We denote this value by $PS$ hereafter.

As shown in Figs. S4 and S5, $PS$ is low when cells are densely adhered to each other, and high when there is a hole in the center.

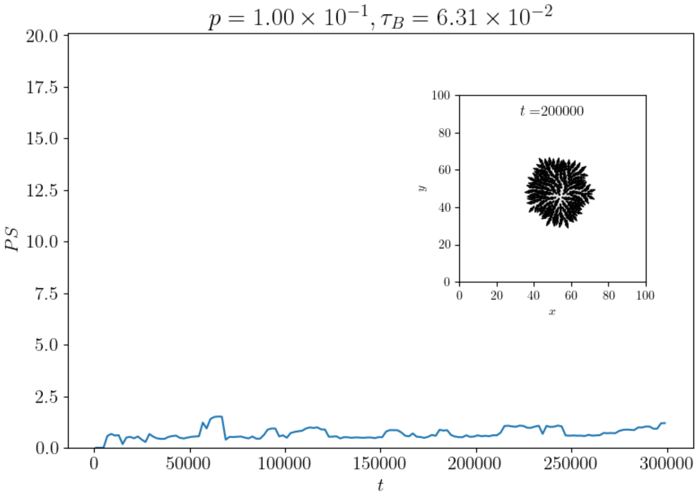


**Fig S4.** Time series of persistence of hole in a single simulation with $p = 1.00 \times 10^{-1}$ and $\tau_B = 6.31 \times 10^{-2}$.

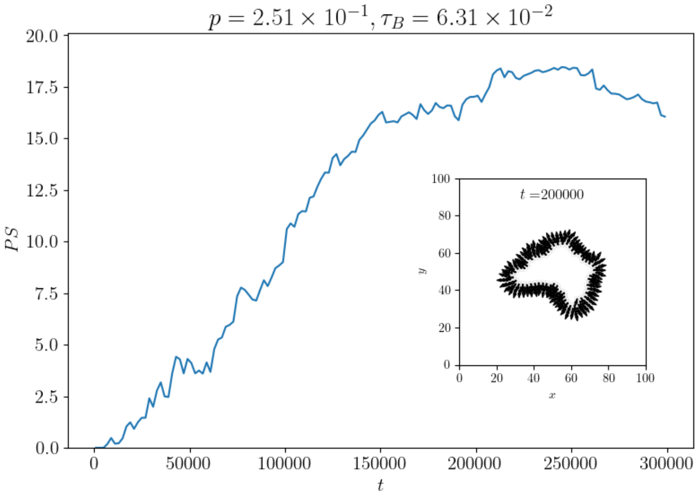


**Fig S5.** A time series of persistence of hole in a single simulation with $p = 2.31 \times 10^{-1}$ and $\tau_B = 6.31 \times 10^{-2}$.

### 3 Formation process of the hole

Furthermore, in the presence of a hole, we would like to consider an index to distinguish between wraparound and inflation in terms of the morphological changes leading to the formation of the hole. Here, we use the cell adjacency network to find the shortest path between any two cells. Among these paths, we calculate the ratio of the length of the longest pair of paths to the straight-line distance in two-dimensional space (Fig. S6).

$$CI = \frac{r_{\text{shortest path}}}{r_{\text{straight}}} \tag{2}$$

We call this “curve index” ($CI$).

By observing the dynamics of the curve index and the presence or absence of a hole, we can evaluate what shape was taken before the hole was formed. If $CI$ decreases sharply when a hole is generated, that is, when $PS$ exceeds a certain value, it indicates that both ends have adhered to each other in a wraparound manner (Fig. S7). If there is no steep change in $CI$ before and after the hole is generated, we consider the hole to have formed by inflation (Fig. S8). Specifically, we defined wraparound as having a relatively large change in the curve index of greater than 3 ($\Delta CI > 3$) at the moment the hole is generated, and inflation as any hole generation other than wraparound [3].

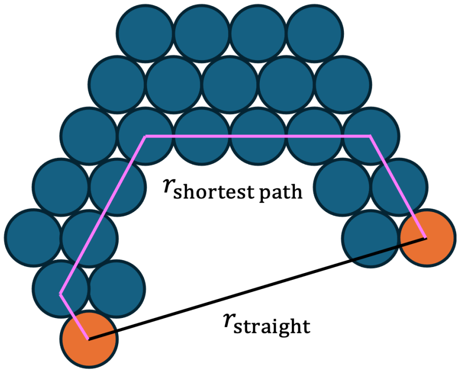


**Fig S6.** Schematic description of the definition of the curve index. For all pairs of cells, the shortest path length on the adjacency network is calculated. The pair with the maximum shortest path length (orange cells in the figure) is used to define $r_{\text{shortest path}}$ (pink path length in the figure). The straight-line distance between these cells in real space is defined as $r_{\text{straight}}$. The curve index is then defined as $r_{\text{shortest path}}/r_{\text{straight}}$.

### 4 Number of layers

The number of layers is also defined as follows.

Here, we consider the number of cells stacked in the direction of polarity for a given cell (Fig. S9). First, we draw a straight line in the direction of polarity with the position of each cell as the center. Then, we count the number of cells that intersect this line and have an inner product with the polarity greater than or equal to $\sqrt{3}/2$. We define this number as the number of layers.

Examples are shown below for the cases of $p = 2.51 \times 10^{-1}$ and $\tau_B = 1.58 \times 10^{-2}$ (Fig. S10) and $p = 3.98 \times 10^{-1}$ and $\tau_B = 1.58 \times 10^{-2}$ (Fig. S11).

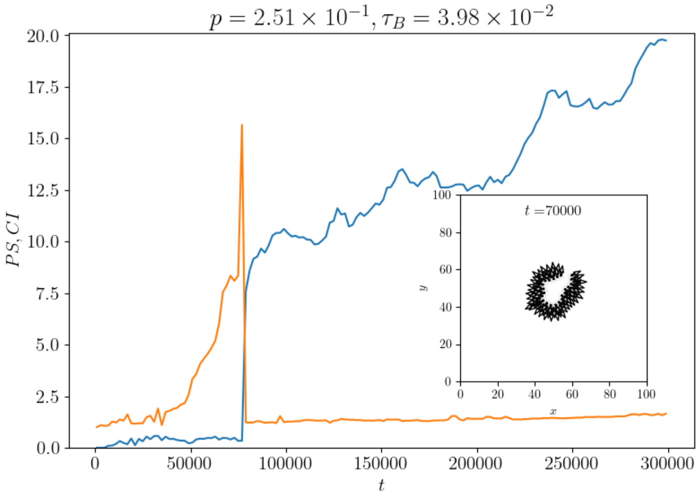


**Fig S7.** Time series of curve index and persistence in a single simulation with $p = 2.51 \times 10^{-1}$ and $\tau_B = 3.98 \times 10^{-2}$.

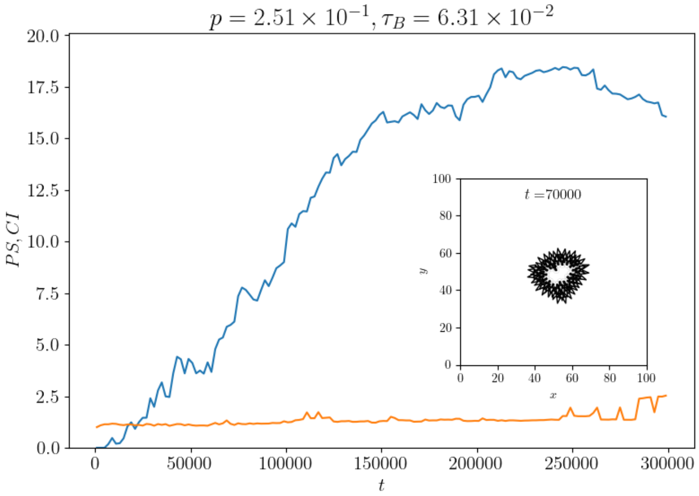


**Fig S8.** Time series of curve index and persistence in a single simulation with $p = 2.51 \times 10^{-1}$ and $\tau_B = 6.31 \times 10^{-2}$.

### 5 Mean field of polarity

Finally, we define the mean field of polarity. This is simply the average of the polarity vectors of all cells.

$$\boldsymbol{P} = \frac{1}{N}\sum_{i}^{N}\boldsymbol{p}_i \tag{3}$$

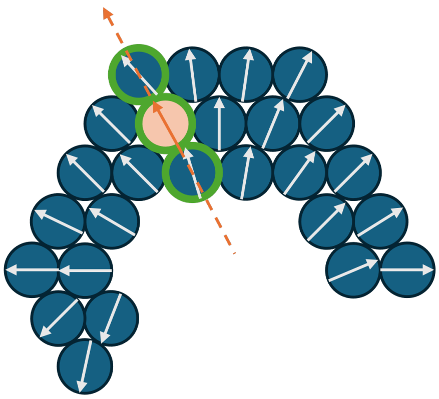

**Fig S9.** Schematic description of the definition of “the number of layers” of the system. For a given cell (light orange cell in the figure), the number of cells that lie along the extension of its polarity direction and have a cosine similarity of polarity greater than or equal to $\sqrt{3}/2$ (cells with green borders in the figure) is defined as the number of layers for that cell. In this figure, the number of layers for the orange cell is 3. This calculation is performed for all cells, and the average is defined as the number of layers in the system.

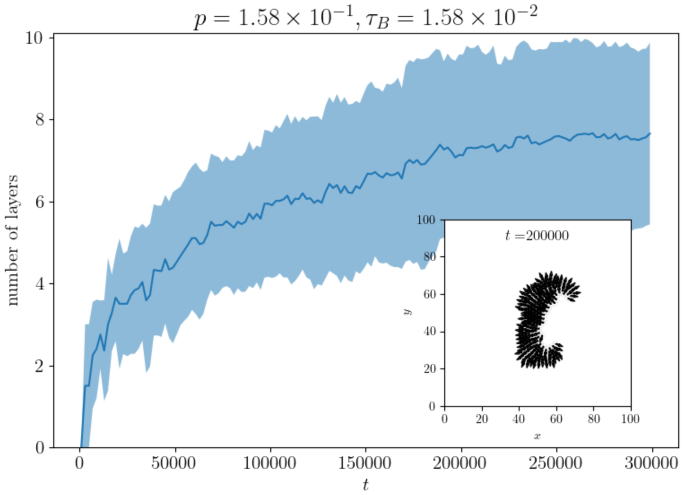


**Fig S10.** Time series of the number of layers in a single simulation with $p = 2.51 \times 10^{-1}$ and $\tau_B = 1.58 \times 10^{-2}$. The line represents the number of layers in the system. The shaded area represents the standard deviation of the number of layers defined for each cell.

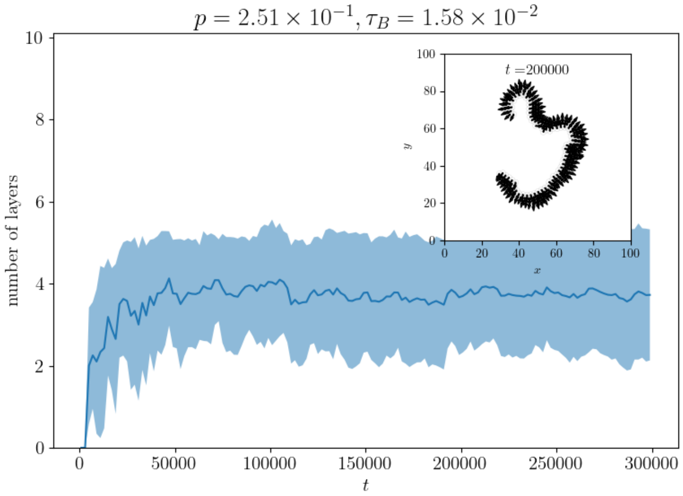


**Fig S11.** Time series of the number of layers in a single simulation with $p = 3.98 \times 10^{-1}$ and $\tau_B = 1.58 \times 10^{-2}$.

### S2 Text: Model for Fig.11

In this modified model, in addition to the dynamics of cell positions and polarity, we consider the following two time evolutions:

- Diffusion of the signal in the field
- Time evolution of the strength of cell polarity ($p$)

The dynamics of the signal field $s(\boldsymbol{r}, t)$ are given by the following reaction-diffusion equation if the position $\boldsymbol{r}$ is not occupied by any cell.

$$\frac{\partial s(\boldsymbol{r},t)}{\partial t} = D_s \nabla^2 s(\boldsymbol{r},t) - \sum_i k_i, \tag{4}$$

where

$$k_i = \begin{cases} c\max(s(\boldsymbol{r},t) - S_i(t), 0) & \text{if } ||\boldsymbol{r} - \boldsymbol{r}_i|| < r_0 \\ 0 & \text{otherwise.} \end{cases} \tag{5}$$

$D_s$ is the diffusion coefficient of the signal. $c$ is a constant that determines the uptake rate of the signal by the cell. $r_0$ is the threshold for the range within which a cell can uptake the signal from the surrounding region. $S_i$ is the amount of signal that cell $i$ has internally. $S_i$ is updated as follows.

$$\frac{\mathrm{d}S_i}{\mathrm{d}t} = \int_{||\boldsymbol{r}-\boldsymbol{r}_i||<r_0} k_i \mathrm{d}\boldsymbol{r} \tag{6}$$

If $\boldsymbol{r}$ is occupied by cell $i$ (i.e., $||\boldsymbol{r} - \boldsymbol{r}_i|| < 1$), the signal field $s(\boldsymbol{r}, t)$ is not updated, and the dynamics is given as follows.

$$\frac{\partial s(\boldsymbol{r},t)}{\partial t} = 0 \tag{7}$$

$$s(\boldsymbol{r},t) = 0. \tag{8}$$

These equations mean that the signal diffuses through the field without penetrating the cells, and is taken up by the cells when it is in their vicinity.

Then, we consider that the strength of cell polarity $p_i$ changes according to the amount of internal signal $S_i$. Here, we assume that the strength of polarity $p_i$ for cell $i$ is determined by the following dynamics.

$$\frac{\mathrm{d}p_i}{\mathrm{d}t} = \tau_p(\sigma(S_i - \theta, \beta) - p_i), \tag{9}$$

where $\tau_p$ is the time constant of the polarity dynamic, $\theta$ is the threshold of the signal for the polarity to be activated, and $\sigma(x, \beta)$ is the sigmoid function defined as follows.

$$\sigma(x,\beta) = \frac{1}{1 + \exp(-\beta x)}. \tag{10}$$

Eq. (9) means that the polarity of cell $i$ increases when the signal $s_i$ is above the threshold $\theta$, and decreases when it is below the threshold.

The initial value of the signal field is $s(\boldsymbol{r}, t = 0) = 2.0$, and for the first cell (cell 1), the initial values are $S_1(t = 0) = 2.0$ and $p_1(t = 0) = 0$. Additionally, when a cell division occurs, the daughter cells inherit the same values of $S$ and $p$ as the mother cell. The parameters used in this model are shown in Table S1.

| Parameter | Value | Description |
|---|---|---|
| $D_s$ | 20 | Diffusion coefficient of the signal |
| $c$ | 0.007 | Uptake rate of the signal by the cell |
| $r_0$ | 1.0 | Threshold for how far a cell can uptake the signal |
| $\tau_p$ | 5.0 | Time constant of the polarity dynamic |
| $\theta$ | 0.5 | Threshold of the signal for the polarity to be activated |
| $\beta$ | 100 | Parameter of the sigmoid function |

**Table S1.** Parameters used in the simulations shown in Fig. 11.

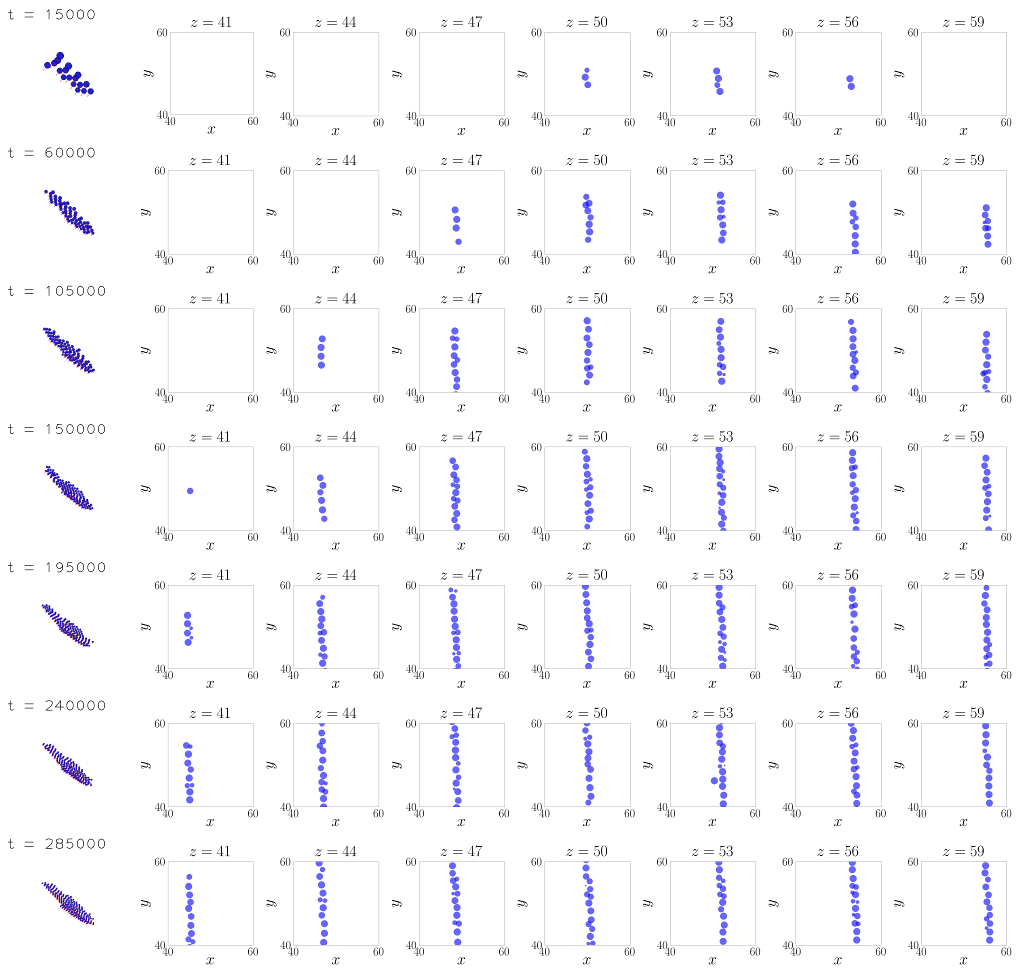


**Fig S12.** 3D version of monolayer elongation. $p = 1, \tau_B = 0.01$. The vertical axis represents time ($t = 15000, 60000, 105000, 150000, 195000, 240000$, and $285000$). The leftmost figure displays a view from an angle in three dimensions. The red arrows indicate the polarities for each cell. The figures arranged horizontally are the $xy$-planes when the $z$-axis is cut at $z = 41, 44, 47, 50, 53, 56$, and $59$. Here, the radius of the cells is assumed to be 1. $\tau_{\text{div}} = 1000.$ is used in the simulations shown in Figs. S12-S18.

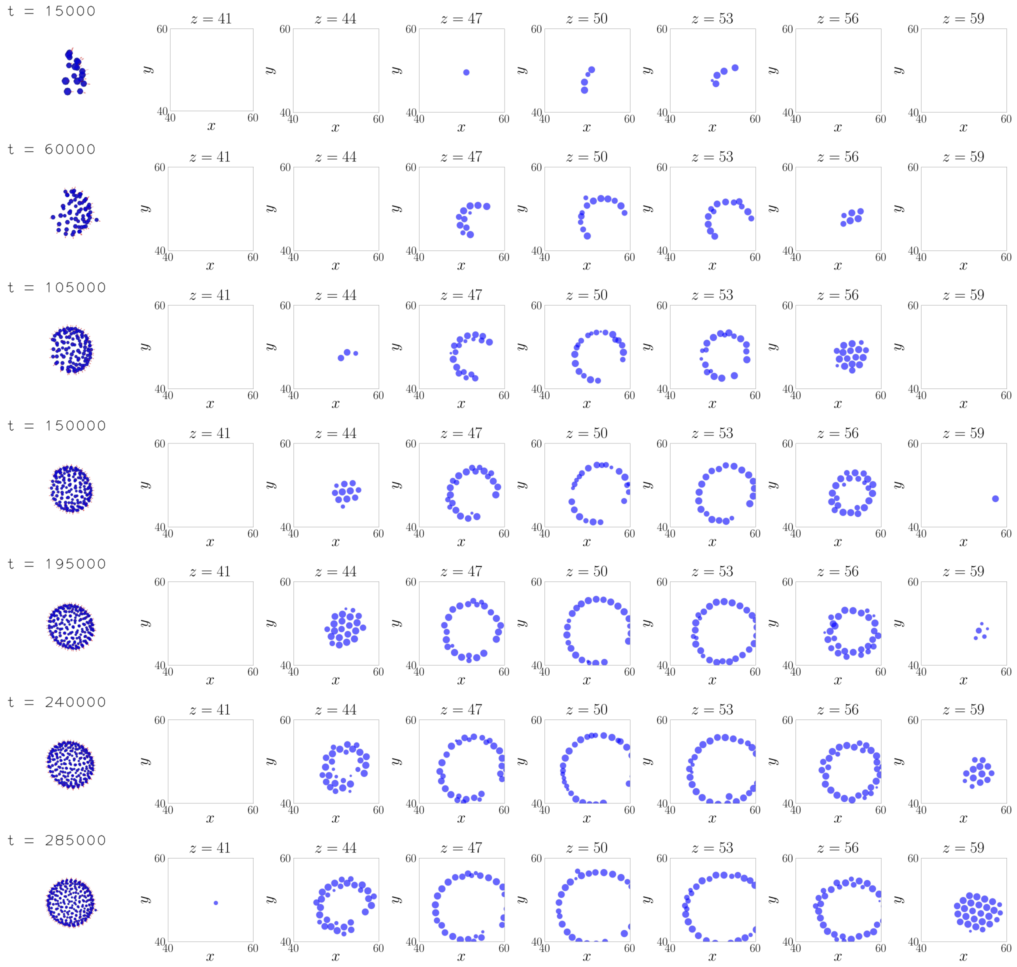


**Fig S13.** 3D version of monolayer wraparound. $p = 1, \tau_B = 1$. For the following figures S13-S18, the same display method is adopted as Fig. S12.

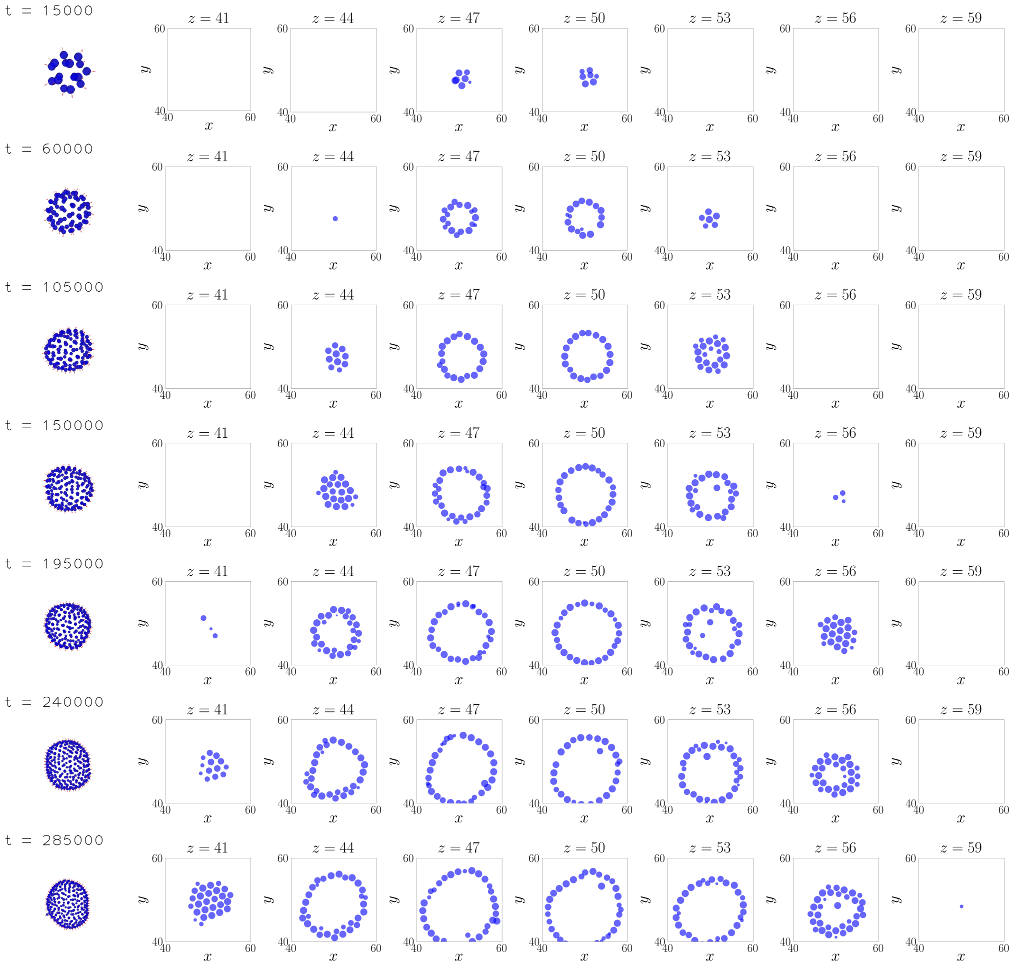


**Fig S14.** 3D version of monolayer inflation. $p = 1, \tau_B = 10$.

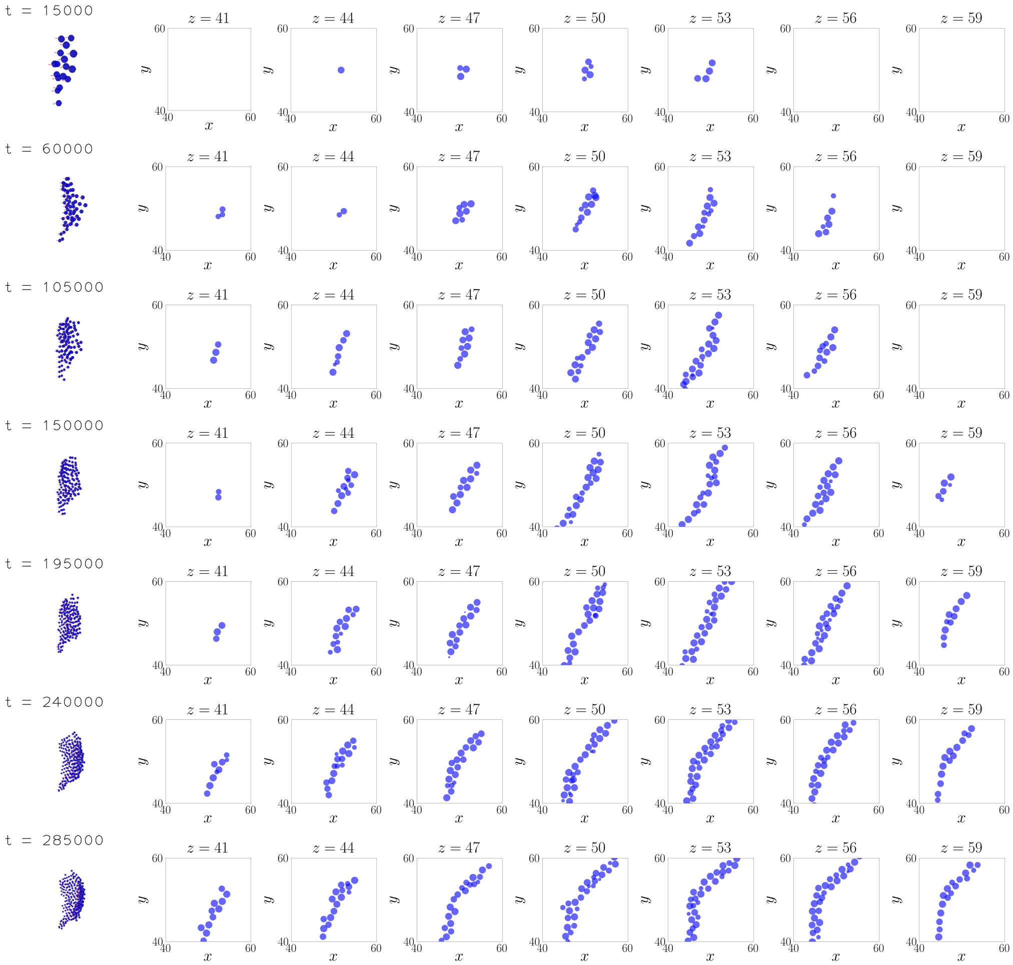


**Fig S15.** 3D version of multilayer elongation. $p = 0.25, \tau_B = 0.01$.

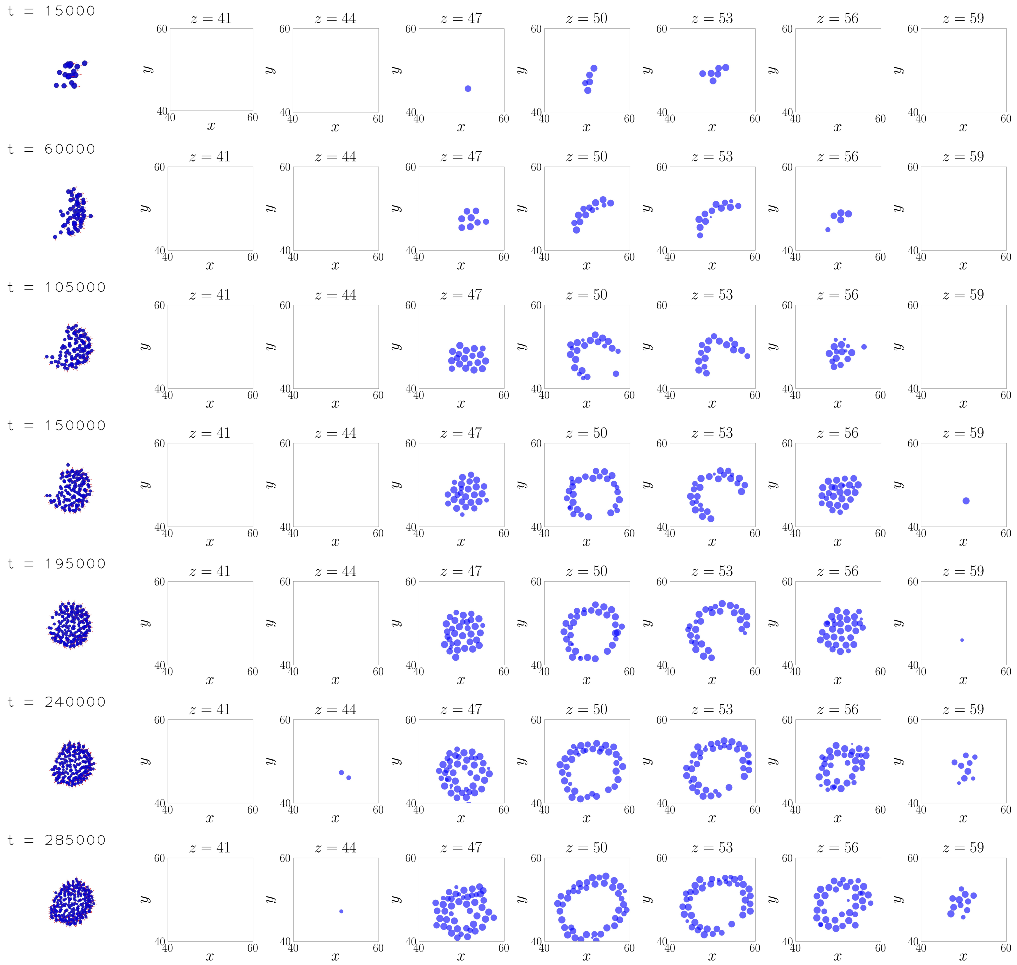


**Fig S16.** 3D version of multilayer wraparound. $p = 0.25, \tau_B = 2.5$.

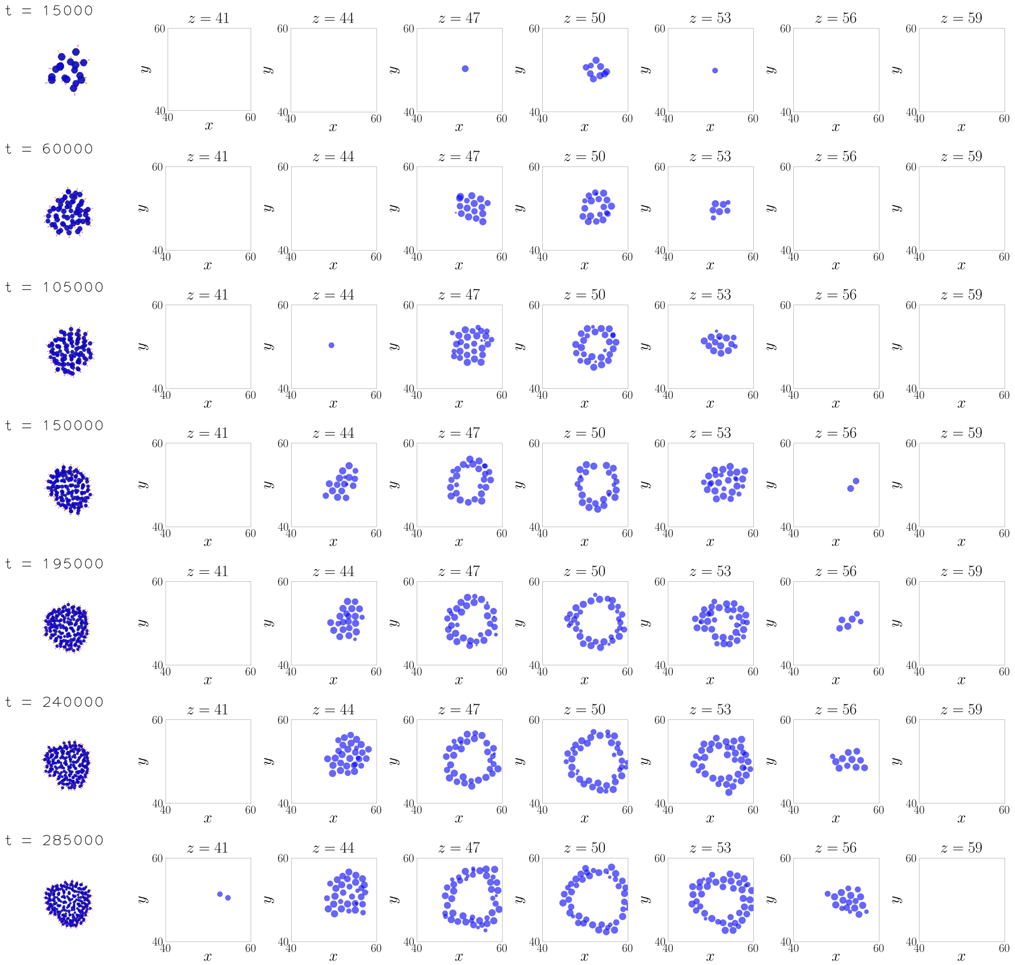


**Fig S17.** 3D version of multilayer inflation. $p = 0.25, \tau_B = 10$.

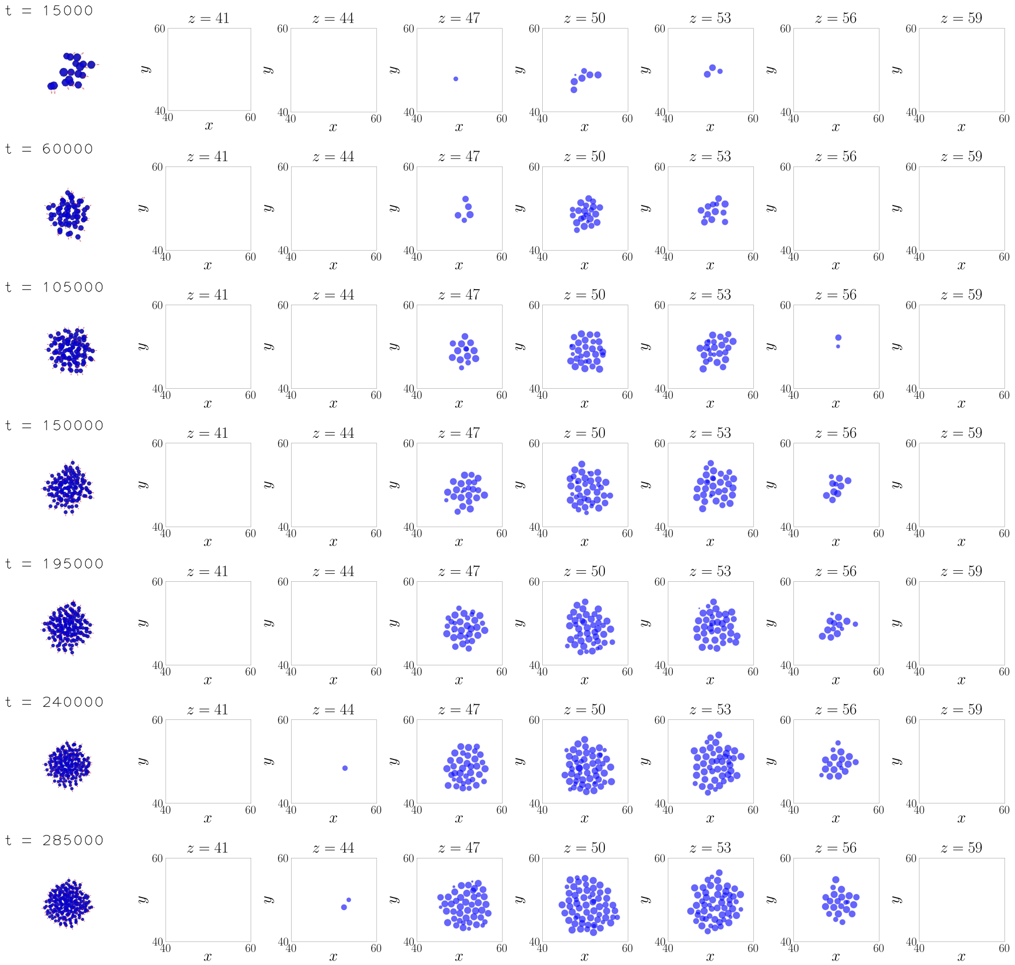


**Fig S18.** 3D version of radial mass. $p = 0.1, \tau_B = 10$.

### References


1. Zomorodian A, Carlsson G. Computing persistent homology. In: Proceedings of the twentieth annual symposium on Computational geometry; 2004. p. 347–356.

2. In practice, to account for some noise, we use $R_{\mathrm{cell}} = 1.25$ for the determination, instead of $R_{\mathrm{cell}} = 1$.

3. Regarding this threshold, values around 1 to 3 do not substantially affect the classification.
